# Evolution from mixed to fixed handedness in mirror-image flowers: insights from adaptive dynamics

**DOI:** 10.1101/2023.07.08.548200

**Authors:** Marco Saltini, Spencer C.H. Barrett, Eva E. Deinum

**Affiliations:** Mathematical & Statistical Methods, Plant Science Group, Wageningen University, 6708 PB Wageningen, The Netherlands; Department of Ecology and Evolutionary Biology, University of Toronto, Toronto, Ontario, Canada M5S 3B2

**Keywords:** adaptive dynamics, enantiostyly, evolutionary transitions, floral asymmetry, pollination, stylar polymorphism

## Abstract

Mirror-image flowers (enantiostyly) involve a form of sexual asymmetry in which a flower’s style is deflected either to the left or right side, with a pollinating anther orientated in the opposite direction. This curious floral polymorphism, which was known but not studied by Charles Darwin, occurs in at least 11 unrelated angiosperm families and represents a striking example of adaptive convergence in form and function associated with cross-pollination by insects. In several lineages, dimorphic enantiostyly (one stylar orientation per plant, both forms occurring within populations) has evolved from monomorphic enantiostyly, in which all plants can produce both style orientations. We use a modelling approach to investigate the emergence of dimorphic enantiostyly from monomorphic enantiostyly under gradual evolution. We show using adaptive dynamics that depending on the balance between inbreeding depression following geitonogamy, pollination efficiency and plant density, dimorphism can evolve from an ancestral monomorphic population. In general, the newly emergent dimorphic population is stable against invasion of a monomorphic mutant. However, our model predicts that under certain ecological conditions, e.g., a decline of pollinators, dimorphic enantiostyly may revert to a monomorphic state. We demonstrate using population genetics simulations that the observed evolutionary transitions are possible assuming a plausible genetic architecture.

## Introduction

Flowering plants possess extraordinary structural variation in floral morphology, particularly in animal-pollinated lineages. Much of this diversity is functionally associated with the pollination biology (Grant and Grant, 1965; Harder and Johnson, 2009; Smith et al., 2009) and mating systems of populations (Barrett and Harder, 2017; Goodwillie et al., 2010; Lloyd and Schoen, 1992). A striking feature of the floral biology of angiosperms is the evolutionary flexibility of reproductive traits which frequently differ between closely related taxa (Barrett et al., 1996; Stebbins, 1974), and also among populations within species (Herlihy and Eckert, 2007; Yuan et al., 2023). This variation is commonly the signature of both past and contemporary evolutionary transitions in floral strategies and mating patterns in response to changing ecological and demographic conditions (reviewed in Barrett, 2008; Pannell and Jordan, 2022). Although most floral traits associated with pollination and mating exhibit continuous patterns of phenotypic variation, caused by interactions between quantitative inheritance and environmental factors (e.g., Ashman, 1999; Caruso, 2004; Fishman et al., 2002), species in numerous families display sexually polymorphic traits with discrete (discontinuous) variation (reviewed in Barrett, 1992; Geber et al., 1999). Commencing with Darwin’s seminal book “Form of Flowers” (Darwin, 1877), polymorphic sexual systems have held a special fascination for evolutionary biologists interested in the evolution and functional significance of reproductive adaptations.

Darwin devoted much of “Forms of Flowers” to the floral polymorphism heterostyly in which populations are composed of two or three floral morphs differing reciprocally in stigma and anther height (Darwin, 1877) (see also Table 1 for a glossary of terms). This polymorphism is reported from 28 angiosperm families (Barrett et al., 2000; Lloyd and Webb, 1992*a*). Darwin interpreted this floral arrangement as a mechanism that functions to promote animal-mediated cross-pollination between floral morphs. Experimental studies on pollen transport and mating in heterostylous populations have largely confirmed Darwin’s cross-promotion hypothesis (Barrett and Shore, 2008; Ganders, 1979; Lloyd and Webb, 1992*a*). After reading an article in The American Naturalist by Todd (1882) on *Solanum rostratum*, Darwin became intrigued by another type of animal-pollinated stylar polymorphism – enantiostyly (mirror-image flowers) – which also exhibits reciprocal sex-organ positioning but involves left- and right-deflected styles. Darwin wrote to Todd requesting seeds of *S. rostratum* so that he could investigate the function of mirror-image flowers but died nine days after writing the letter to Todd (Darwin, 1882, reprinted in 1945; Jesson and Barrett, 2005). Interest in these curious left-right stylar asymmetries remained largely dormant - until relatively recently (reviewed in Barrett and Fairnie, 2024), when workers began to investigate various unrelated enantiostylous taxa (e.g., Bowers, 1975; Dulberger, 1981; Fenster, 1995; Graham and Barrett, 1995; Jesson and Barrett, 2003; Mora-Carrera et al., 2019; Ornduff and Dulberger, 1978; Richman and Venable, 2018; Wang, 1995). However, in contrast to the extensive ecological and genetic research conducted on heterostylous taxa over the past century (reviewed in Barrett, 1992; Ganders, 1979; Lloyd and Webb, 1992*a*,*b*), the evolutionary origins and adaptive significance of enantiostyly remain poorly understood.

**Table 1:** Glossary of terms related to plant reproduction used in this article.

| Term | Description |
| --- | --- |
| Dioecy | A sexual system in which populations are composed of female and male plants. |
| Enantiostyly | A floral polymorphism in which styles are deflected either to the left- or right-side of a flower. In monomorphic enantiostyly, both flower types can occur on a plant (“mixed handedness”), whereas in dimorphic enantiostyly plants produce either left- or right-handed flowers (“fixed handedness”). |
| Floral display | The number of flowers open on a plant at one time (e.g., daily display size). |
| Floral longevity | The functioning life span of an individual flower. |
| Geitonogamy | Self-pollination resulting from pollen transfer between flowers on a plant that causes self-fertilization (selfing). |
| Herkogamy | The spatial separation of dehiscing anthers and receptive stigmas within a flower. |
| Heterostyly | A sexual polymorphism in which populations are composed of two (distyly) or three (tristyly) floral morphs differing reciprocally in stigma and anther heights. |
| Inbreeding depression | The reduction in viability and/or fertility of inbred offspring compared to those from mating between unrelated individuals. A common form of inbreeding in plants is self-fertilization (selfing). |
| Mating system | The mode of transmission of genes from one generation to the next through sexual reproduction. |
| Monoecy | A hermaphroditic sexual system in which plants in a population produce separate female and male flowers. |
| Pollen discounting | A loss in outcrossed siring success caused by self-pollination. |
| Pollination | The transfer of pollen between flowers by various agents of pollen dispersal including animals, wind and water. |
| Stylar dimorphism | The occurrence of two floral morphs within a population that differ in style orientation (e.g., dimorphic enantiostyly) or length (e.g., distyly). |

Reluctance to tackle the problem of how and why mirror-image flowers evolved may be because of a puzzling feature of the floral polymorphism. In contrast to heterostyly and other stylar polymorphisms (summarized in Barrett, 2010, Table 2), in which all flowers borne by an individual possess the same stylar condition and populations are reproductively subdivided into distinct floral morphs, mirror-image flowers exist in two fundamentally different forms (see Fig. 2 in Barrett, 2002). In monomorphic enantiostyly, individual plants produce a mixture of both left- and right-handed flowers (hereafter L- and R-flowers), whereas in species with dimorphic enantiostyly, plants are fixed for stylar orientation (hereafter L- and R-plants). Thus, monomorphic enantiostyly involves somatic floral polymorphism and only dimorphic enantiostyly resembles other stylar polymorphisms in being a true genetic polymorphism. In *Heteranthera missourienisis* (previously *H. multiflora*), the only dimorphic enantiostylous species in which the genetic basis of handedness has been investigated, right-handedness is governed by a dominant allele at a single diallelic Mendelian locus with left-handed plants homozygous recessive (Jesson and Barrett, 2002*a*).

**Table 2:** Model parameters (A) before and (B) after non-dimensionalisation.

| A |  |  |
| --- | --- | --- |
| Parameter | Description | Unit |
| $c$ | Pollen-to-seed conversion coefficient | - |
| $f$ | Number of flowers in daily display | - |
| $\delta$ | Fitness cost of inbreeding depression following geitonogamy | - |
| $\gamma$ | Efficiency reduction for pollen transfer between flowers of the same stylar orientation | - |
| $h$ | Handling time | day |
| $r_p$ | Pollination rate | day<sup>-1</sup> |
| $m_0$ | Plant base mortality rate | day<sup>-1</sup> |
| $S_I$ | Foraging area of pollinators | m<sup>2</sup> |
| $S_C$ | Plant competition area | m<sup>2</sup> |
| $x$ | Fraction of right-handed flowers on an individual | - |
| $N_x$ | Density of plants with phenotypic trait $x$ | m<sup>-2</sup> |

| B |  |  |
| --- | --- | --- |
| Parameter | Description | Expression |
| $\varphi$ | Reproductive rate | $cr_p f^2 / m_0$ |
| $\chi$ | Handling time | $m_0 h / c$ |
| $\sigma$ | Competition coefficient | $S_C / S_I$ |
| $\nu_x$ | Density of plants with phenotypic trait $x$ | $S_I N_x$ |

Comparative analysis of several enantiostylous families (e.g. Solanaceae, Gesneriaceae, Haemodoraceae, Pontederiaceae) supports an evolutionary progression from a straight-styled ancestral condition to monomorphic enantiostyly, with the evolution of left-right stylar asymmetry within plants followed by rare transitions to dimorphic enantiostyly (reviewed in Jesson and Barrett, 2003) restricted to a few monocotyledonous lineages (Graham and Barrett, 1995). Thus, the vast majority of enantiostylous species distributed across at least 11 families possess monomorphic enantiostyly. This uneven distribution of the two enantiostylous conditions is puzzling for two reasons. First, no new developmental machinery appears to be required to produce stylar dimorphism as this already exists in the monomorphic ancestral state. Second, experimental studies on the function of the two forms of enantiostyly demonstrate clear mating benefits to stylar dimorphism because it limits selfing by geitonogamous pollen transfer (Barrett et al., 2000; Jesson and Barrett, 2002*b*, 2005). Because pollen deposition on the right- or left-side of the bodies of foraging bees by pollinating anthers favors cross-pollination between flowers of different stylar orientation rather than between flowers of the same stylar deflection, rates of geitonogamy are significantly higher for monomorphic than dimorphic enantiostyly, although the former condition does reduce geitonogamy in comparison with the straight-styled ancestral condition (Jesson and Barrett, 2002*b*, 2005). Geitonogamy is generally recognized as a non-adaptive cost of floral display size as it can result in inbreeding depression and pollen discounting (Harder and Barrett, 1995; Lloyd, 1992, although see Mora-Carrera et al. (2019) for an adaptive explanation for geitonogamy in a monomorphically enantiostylous species).

A variety of hypotheses has been proposed to explain the overall rarity of dimorphic enantiostyly involving potential evolutionary constraints on transitions from monomorphism to dimorphism (Jesson et al., 2003*a,b*). These include weak selection for fixed handedness in populations less susceptible to geitonogamy owing to their small daily floral display sizes; for example, in monomorphic species in which only a single flower is displayed each day (Barrett et al., 2000). A lack of heritable genetic variation for the direction of asymmetry in monomorphic populations, and developmental-genetic constraints because of the absence of genes with appropriate positional information to distinguish left from right (Jesson and Barrett, 2002*a*; Jesson et al., 2003*b*), especially in species with otherwise radially symmetric flowers (see Coen and Meyerowitz, 1991; Luo et al., 1996).

Insights into the possible conditions required for evolutionary transitions between monomorphic and dimorphic enantiostyly can be obtained by theoretical approaches. Jesson et al. (2003*a*) approached this problem using phenotypic selection models. They did so in two steps: first, by investigating how monomorphic enantiostyly may have evolved from a straight-styled ancestor through gradual selection of increased stigma-anther separation (herkogamy). Then, how dimorphic enantiostyly could evolve from monomorphic enantiostyly through the successive invasion of two variants with fixed handedness (either L- or R-plants). Although Jesson et al. (2003*a*) also hypothesized that dimorphic enantiostyly could have evolved from monomorphic enantiostyly through disruptive selection on the proportion of L- and R-flowers within a plant they did not explore this hypothesis theoretically.

Here, we investigate the gradual adaptive evolution of the proportion of L- and R-flowers within individuals of a monomorphic enantiostylous population using the adaptive dynamics framework (Dieckmann and Law, 1996; Geritz et al., 1998; Metz et al., 1996; van Cleve, 2023) to derive the fitness of invading variants directly from population dynamic considerations, complemented with population genetic simulations. We focus primarily on the evolution of dimorphic enantiostyly from monomorphic enantiostyly as, to our knowledge, there is no comparative evidence of the reverse transition in angiosperms, although we do consider when this might occur in our models. Our analyses address the following specific questions: (i) what ecological factors influencing population dynamics lead to the emergence of dimorphic enantiostyly from monomorphic enantiostyly via evolutionary branching and gradual adaptive evolution? (ii) Once established, what ecological and demographic conditions maintain the newly arisen stylar dimorphism resulting in evolutionary stability?

Our analyses link ecological factors to the evolution of the ratios of L- and R-flowers within individuals. By doing so, we can study what environmental conditions promote the occurrence of evolutionary branching and the emergence of stylar dimorphism from monomorphism. With this approach, we explicitly derive the invasion fitness of a variant carrying a mutation that influences the ratio of L- and R-flowers from the ecological interactions of such a variant with individuals with the resident phenotypic trait. This readily reveals that such interactions are a source of negative frequency-dependent selection, the principal selective mechanism that maintains sexual polymorphisms. In line with similar models of the evolution of traits influencing plant reproduction (Cheptou and Mathias, 2001; de Jong and Geritz, 2001; de Jong et al., 1999), we assume that evolution proceeds by rare mutations of small phenotypic effects governing the proportion of L- and R-flowers within individuals. We show that under particular environmental conditions geitonogamous selfing resulting from pollen transfer between flowers on the same plant drives the evolution of dimorphic enantiostyly from monomorphic enantiostyly. Furthermore, we also examine how changes in the environment might under certain conditions cause the breakdown of dimorphic enantiostyly and restoration of a monomorphic state. Finally, we provide proof-of-principle that the transition from monomorphic to dimorphic is possible with a pattern of inheritance similar to that of *H. missouriensis* (Jesson and Barrett, 2002*a*).

## 2 The model

We study the adaptive evolution of a population of enantiostylous plants, where the evolving phenotypic trait *x* is the average fraction of R-flowers compared to the total average number of flowers *f* displayed daily on a plant, and 1 − *x* is the fraction of L-flowers. In the following sections, we introduce an ecological model that describes the dynamics of a monomorphic enantiostylous population and derive an explicit analytical expression for the interaction between plants with different phenotypic traits. This interaction, mediated by the movement of pollen by pollinators, controls the adaptive evolution of the phenotypic trait *x*. In this way, we create a link between the dynamics of the population and the evolutionary dynamics of the trait *x*. Following this, we extend the ecological model describing the dynamics of a monomorphic population to include scenarios involving multiple interacting phenotypes. With this extension, we investigate co-evolutionary dynamics of multiple phenotypes emerging from phenotypic diversification. The relevant model parameters are listed in Table 2A.

### 2.1 Ecological dynamics

#### 2.1.1 Pollen transfer between plants

We introduce the pollination rate *r*_*p*_ as the rate at which pollen is transferred between flowers. The efficiency of pollen transfer between flowers with the same or the opposite stylar orientation depends on morphological and behavioural traits of the pollinators. For example, it has been demonstrated that in *Wachendorfia paniculata*, sufficiently large pollinators promote pollination between flowers of opposite stylar orientations, whereas smaller pollinators transfer pollen between flowers of the same or different stylar orientation with more comparable efficiency (Minnaar and Anderson, 2021). Therefore, we assume that the pollination rate *r*_*p*_ is maximum when pollen is transferred between flowers of the opposite stylar orientation, whereas *r*_*p*_ is multiplied by a factor 1 − *γ*, with *γ* ranging from 0 to 1, when pollen is transferred between flowers of the same stylar deflection, see Figure 1A. By doing so, we account for the potentially lower efficiency of large pollinators to move pollen between flowers of the same stylar orientation. The total amount of pollen dispersed from one plant to another is therefore dependent on the number of L- and R-flowers on each plant. Let *x*_1_ and *x*_2_ be the fraction of R-flowers of two plants with *f* flowers each displayed daily. Therefore, the two plants show, respectively, *fx*_1_ and *fx*_2_ R-flowers in the daily display. The rate at which pollen is received by R-flowers of the first plant from L-flowers of the second plant is *r*_*p*_*fx*_1_*f*(1 − *x*_2_). Similarly, *r*_*p*_*f*(1 − *x*_1_)*fx*_2_ is the rate at which pollen from R-flowers of the second plant is received by L-flowers of the first. Then, the rate at which pollen is transported by pollinators between flowers of the same stylar orientation is: *r*_*p*_(1 − *γ*)*fx*_1_*fx*_2_ and *r*_*p*_(1 − *γ*)*f*(1 − *x*_1_)*f*(1 − *x*_2_).

**Figure 1:**
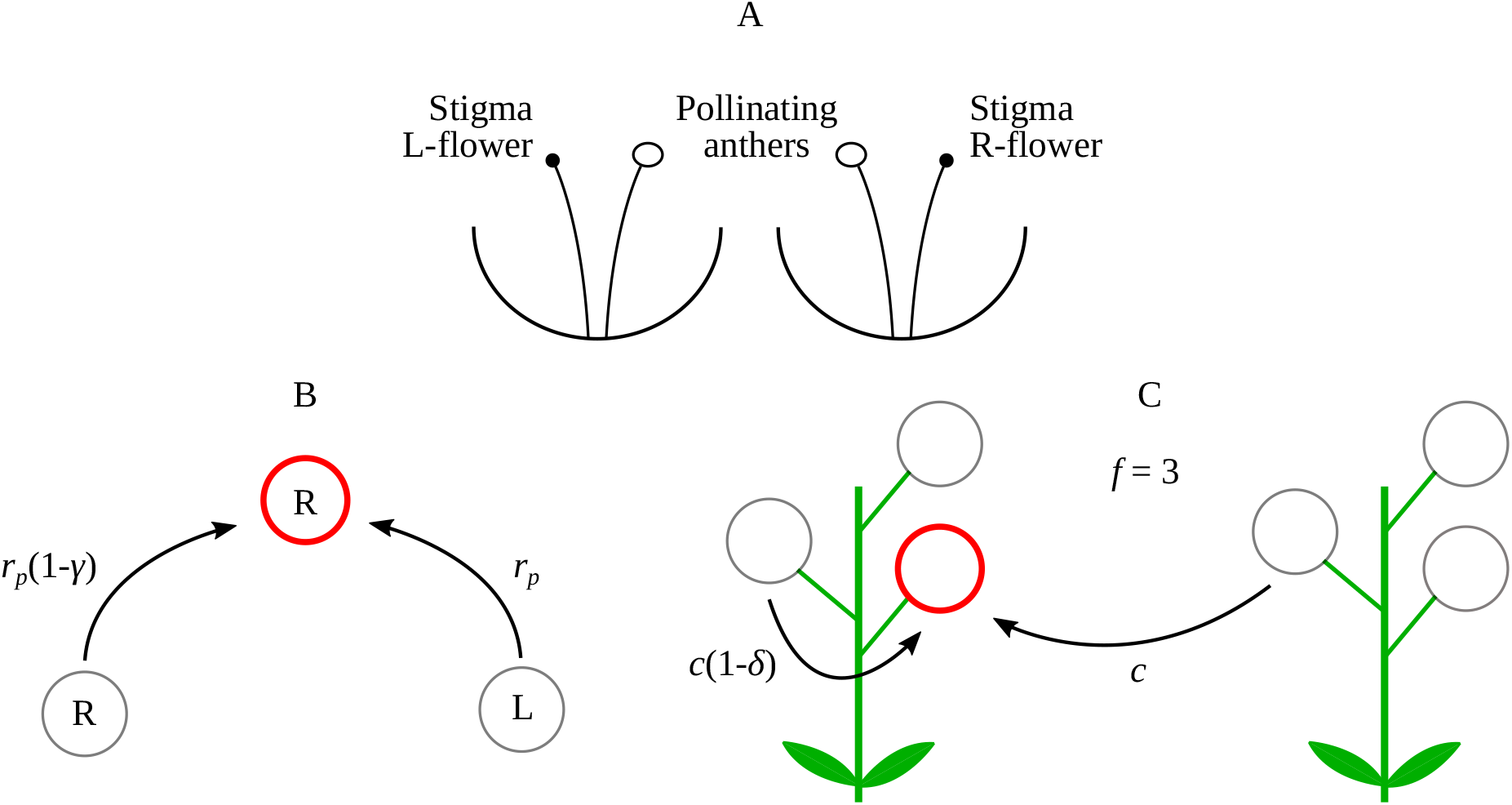
Schematic of the model. (A) Stylised representation of an L-flower and an R-flower in reciprocal enantiostyly. (B) Schematics of hypothetical pollen dispersal among flowers of the same and different stylar orientations of an enantiostylous species. The flower with opposite handedness transfers pollen at the full rate *r*_*p*_, whereas the flower with the same handedness suffers a penalty of 1 − *γ*, compared to pollen movement between flowers of opposite stylar orientation. (C) Schematics of pollen grain-to-seed conversion *c* resulting from fertilization by outcrossing or geitonogamy. In the latter case, the conversion coefficient is reduced by 1 − *δ* compared to outcrossing due to inbreeding depression.

We combine these contributions to give an overall rate of pollen transport: *r*_*p*_*f* ^2^*p*_*γ*_(*x*_1_, *x*_2_), which we hereafter refer to as the “pollination efficiency” and define the “pollination function” as:

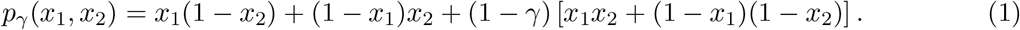

This function is the fraction of pollen deposited on stigmas of any flower of the first plant with respect to the amount of pollen on a pollinator’s body originating from any flower of the second plant. Note that the pollination function is symmetric under interchange of variables, i.e., *p*_*γ*_(*x*_1_, *x*_2_) = *p*_*γ*_(*x*_2_, *x*_1_). Pollinators that are highly specialised for pollen movement between flowers with the opposite stylar orientation have *γ* = 1, and therefore a pollination efficiency *r*_*p*_*f* ^2^*p*_1_(*x*_1_, *x*_2_), with *p*_1_(*x*_1_, *x*_2_) = *x*_1_(1 − *x*_2_) + (1 − *x*_1_)*x*_2_ which is maximum for *x*_1_ = 0 and *x*_2_ = 1 (or *vice versa*). Conversely, if the number of pollen grains dispersed from a flower to another plant of the same or the opposite stylar orientations is roughly equal (*γ* = 0), then the pollination efficiency is *r*_*p*_*f* ^2^*p*_0_(*x*_1_, *x*_2_), with *p*_0_(*x*_1_, *x*_2_) = 1.

#### 2.1.2 Dynamics of a monomorphic population

The population model is based on per-capita reproduction and mortality. Let the trait *x* characterize a plant population with density *N*_*x*_, per-capita reproductive rate *b*(*x*), per-capita mortality rate *m*(*x*). Then, the equation describing the population dynamics of *N*_*x*_ is

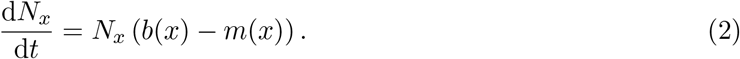

The reproductive rate is split into two terms: seed production following geitonogamous pollen transfer and following outcrossing. In the case of outcrossing, we first define *S*_*I*_ as the foraging area of pollinators. This quantity defines the surface area for plants to exchange pollen with each other. Then, we introduce the conversion coefficient of pollen grains into seeds *c* as the average fraction of one seed produced by a single pollen grain. Thus, the reproductive rate due to outcrossing is proportional to

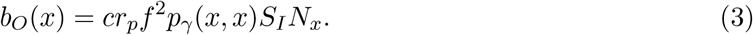

Note that from Equation (3) it follows that a higher density of individuals in an area has the effect of increasing the per-capita reproductive rate by producing more pollen for outcrossing. We hereafter refer to this interaction, which is mediated by pollinators and the intensity of which is controlled by the pollination function, as “facilitation”. In Equation (3), we assume the density of individuals *N*_*x*_ to be high enough to preclude a population from going extinct due to demographic stochasticity.

Similarly, we assume the contribution to the reproductive rate due to geitonogamous pollen transfer to be reduced by a factor 1 − *δ*, which describes the fitness cost of self-fertilization due to inbreeding depression, see Figure 1B. Here, *δ* ranges between 0 (no fitness cost due to geitonogamy) and 1 (complete inbreeding depression). Thus, the reproductive rate following geitonogamy is proportional to

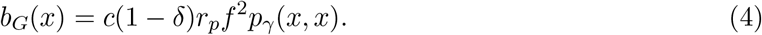

Finally, we include a saturation effect for the overall reproductive rate by assuming a Holling’s type-II functional response to pollen availability. To do so, we introduce a parameter *h* which in the animal literature is typically referred to as handling time (Holling, 1959). In our context, *h* describes the time required for replacing flowers that after fertilization lack receptive ovules. Then, the overall reproductive rate is

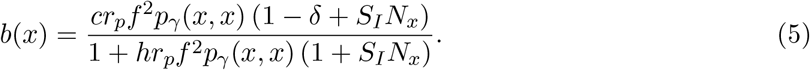

We assume that the per-capita mortality rate of plants is density dependent following

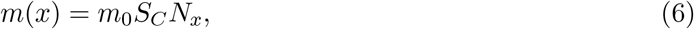

where *m*_0_ is the base mortality rate and *S*_*C*_ is the competition area that represents the range within which different individuals compete with each other for shared abiotic resources (e.g., water, soil nutrients). For mathematical simplicity, we excluded density-independent mortality from our model. Note that this decision may prevent the occurrence of an Allee effect.

#### 2.1.3 Non-dimensionalization

Equation (2) depends on nine distinct parameters (Table 1). To ease the exploration of the parameter space, we reduced the number of parameters through non-dimensionalization. We use the plant base mortality rate *m*_0_ as the unit of time to re-scale the time dependent quantities by defining the non-dimensional time *τ* as *τ* = *m*_0_*t*. We use the foraging area of pollinators *S*_*I*_ to re-scale the length dependent quantities by defining the non-dimensional density of plants *ν*_*x*_ as *ν*_*x*_ = *S*_*I*_*N*_*x*_. We define the non-dimensional reproductive rate *φ* = *cr*_*p*_*f*^2^/*m*_0_, the non-dimensional handling time *χ* = *m*_0_*h/c*, and the non-dimensional competition coefficient *σ* = *S*_*C*_/*S*_*I*_, see Table 2B.

With these, Equation (2) becomes

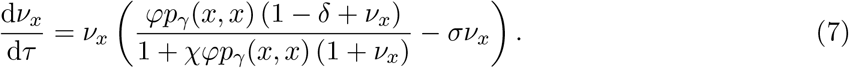

The number of free parameters is now reduced from nine to five.

Equation (7) has two stationary solutions that depend parametrically on the trait value *x*: the trivial *ν*_*x*_ = 0, and 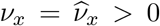 (Equation S1). Because of our assumption of excluding density-independent mortality, *ν*_*x*_ = 0 is a repeller of the population dynamics, while *ν*_*x*_ is an attractor, and will be hereafter used as the equilibrium density for a resident population with trait *x*, see Supplementary Information S1 for detail.

#### 2.1.4 Interaction coefficients

To study the evolution of the trait *x*, it is necessary to understand how different phenotypes interact with each other. In our model, pollen production by individuals facilitates reproduction, which we model by considering the exchange of pollen between all possible pairs of flowers, whereas competitive interactions for shared abiotic resources influence mortality and are thus modelled at the population level.

The facilitative interaction is mediated by the pollinating function defined in Equation (1). Here, unlike the monomorphic case where an offspring had the phenotype of its parent plants, we assumed that the phenotype of an offspring was inherited via ovules with a probability *z*, meaning the offspring shows the same phenotype as the maternal parent, or via pollen with a probability 1 − *z*, i.e., its phenotype is passed by the paternal parent. Therefore, in a population with phenotypic traits *x*_1_, …, *x*_*n*_, the reproductive rate of the phenotype *x*_*i*_ due to outcrossing is proportional to:

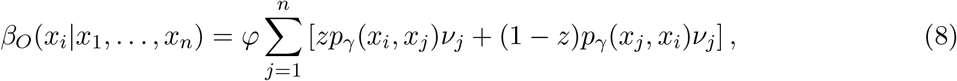

where the summation runs over all possible phenotypes in the population, including both non-geitonogamous pollen transfer between plants of the same phenotypes (when *j* = *i*), and pollen transfer between plants of different phenotypes. Given that *p*_*γ*_(*x*_*i*_, *x*_*j*_) = *p*_*γ*_(*x*_*j*_, *x*_*i*_), the overall reproductive rate of the phenotype *x*_*i*_ is:

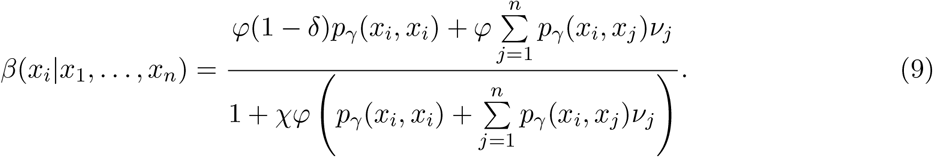

In this equation, the two terms outside of the summations represent geitonogamous pollen transfer. Similar to the reproductive rate, the *per-capita* mortality rate when *n* phenotypes coexist is:

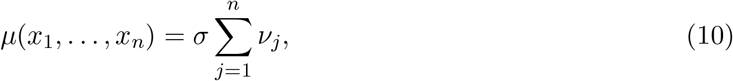

where mortality depends on the competition between plants of both the same (*j* = *i*) and different phenotypes. In Equations (8), (9) and (10), the *ν*_*j*_ term represents the density of plants with phenotype *x*_*j*_.

### 2.2 Evolutionary dynamics

Based on the ecological scenario presented in the previous section, we use invasion analysis (*sensu* Otto and Day, 2007) to study the gradual evolution of the phenotypic trait *x* that characterizes the stylar condition of an initially monomorphic plant population. Specifically, we investigate the conditions allowing an emerging mutant to establish in a population and replace the resident from which it originated. This process, known as the trait substitution sequence (see Champagnat et al., 2006; Dieckmann and Law, 1996) is depicted in Figure 2A,C (black arrows), and produces evolutionary trees as in Figure 2G-I. Whether a mutant can invade a resident population depends on its invasion fitness, defined in the next section.

**Figure 2:**
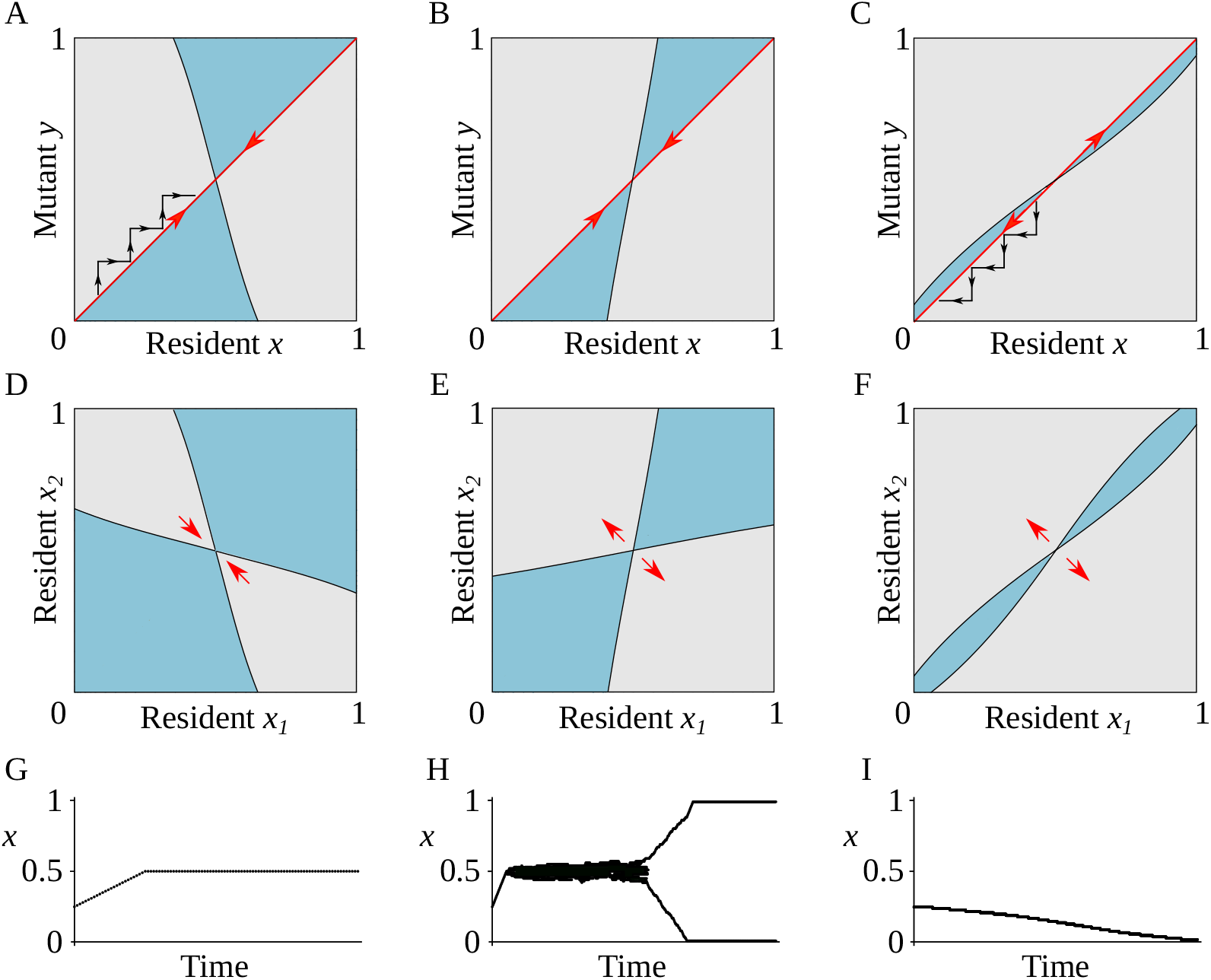
The three possible types of evolutionary behaviour: gradual evolution can result in a population of monomorphic enantiostyly (A,D,G), dimorphic enantiostyly (B,E,H), or a population with uniform direction for stylar orientation (C,F,I). (A-C) Pairwise invasability plots (PIPs). Mutants can invade a resident with trait value *x* if their trait value *y* is in a grey (light) area which occurs when *y* has positive invasion fitness. Blue (darker) regions in the plots correspond to negative invasion fitness and, in those regions, the mutant *y* cannot invade the resident *x*. The red arrows indicate directional evolution (A,B) towards the singular point *x*\* = 1/2 (evolutionary attractor, convergent stable) or (C) away from the singular point *x*\* = 1/2 (evolutionary repeller, not convergent stable), under the assumption of mutations of small phenotypic effect. Evolution proceeds as a trait substitution sequence illustrated by the black arrows in (A) and (C). (D-F) Area of coexistence of a hypothetical dimorphic population, with phenotypic traits *x*_1_ and *x*_2_. Grey (light) areas correspond to regions where *x*_1_ and *x*_2_ can coexist on an ecological time scale. Red arrows represent the direction of (A) convergent and (B,C) disruptive selection in a coevolutionary process involving *x*_1_ and *x*_2_. (G-I) Example of simulated evolutionary trees. (G) After an initial period of directional selection, evolution reaches an end point with a monomorphic population at *x* = 1/2. (H) Following directional evolution, the monomorphic population splits into two subpopulations with phenotypic traits *x* = 0 and *x* = 1. (I) Directional selection drives the monomorphic population away from the singular point and towards *x* = 0. Note that with a different starting point (*x* > 1/2), the evolutionary process would go towards *x* = 1. Parameter values: (A,D,G) *φ* = 10, *χ* = 1, *σ* = 5, *δ* = 0.6, *γ* = 0.8. (B,E,H) *φ* = 10, *χ* = 1, *σ* = 5, *δ* = 0.8, *γ* = 0.5. (C,F,I) *φ* = 15, *χ* = 1, *σ* = 5, *δ* = 0.9, *γ* = 0.2.

#### 2.2.1 The invasion fitness and the trait substitution sequence

A single step of the adaptive dynamic process consists of the introduction of an initially rare mutant with phenotypic trait *y* in a population of residents with trait *x* at its population dynamic equilibrium 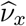 (Equation S1). We focus on gradual adaptive evolution and therefore we assume mutations of small phenotypic effects, i.e., *y* = *x* ± *ε*, with *ε* ≪ 1.

The mutation is initially carried by few individuals and, together with the assumption of high density for the resident population, the probability of pollen transfer among different mutants is negligible. Nor does the mutation cause density dependent effects on the mortality rate of the resident population. Then, the invasion fitness *ρ*(*y*|*x*) of the rare mutant *y* in a resident population that consists of a single trait value *x* is the initial *per-capita* growth rate of the mutant population (Geritz et al., 1998). In our case, we have:

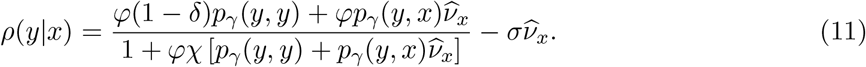

Mutants with positive invasion fitness can invade the resident population, whereas those with negative invasion fitness are unable to do so. Under the adaptive dynamics assumptions, a mutant that can invade will typically replace the resident population and become the new resident, resulting in a trait substitution sequence (Dieckmann and Law, 1996). This occurs when *ρ*(*y*|*x*) > 0 and *ρ*(*x*|*y*) < 0. Through this, the evolving trait climbs the dynamic fitness landscape and evolves to locally maximize its fitness. Instead, if both conditions *ρ*(*y*|*x*) > 0 and *ρ*(*x*|*y*) > 0 hold, the mutant and resident can coexist and evolution results in so-called evolutionary branching and the creation of dimorphism. The latter condition occurs when the resident trait is at a minimum of the invasion fitness function.

Under the adaptive dynamics assumption, the speed of evolution and direction of the trait substitution sequence follows from the selection gradient *s*(*x*) = ∂*ρ/*∂*y*|_*y*=*x*_, defined in Equation (S8). The trait substitution process continues until a singular point of the evolutionary dynamics is reached. In a singular point, the selection gradient vanishes, i.e., *s*(*x*\*) = 0. It can easily be verified that *x*\* = 1/2 is a singular point. Note that *x*\* = 1/2 corresponds to a 1:1 ratio of L- and R-flowers on individual plants, as often observed in natural populations with monomorphic enantiostyly (Jesson and Barrett, 2003). It may be that under specific conditions, the equation *s*(*x*\*) = 0 can also be satisfied for a values of *x*\* other than *x*\* = 1/2, see Supplementary Information S4. However, with our exploration of the parameters space we could not find parameters combinations such that solutions other than *x*\* = 1/2 exist. Therefore, if they exist, they would likely be idiosyncratic and, as such, largely irrelevant for the question of how dimorphic enantiostyly could gradually evolve from monomorphic enantiostyly.

How evolution proceeds at a singular point is determined by the signs of the functions *M*(*x*\*) and *E*(*x*\*) defined, respectively, as

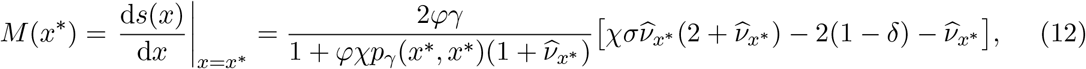

and

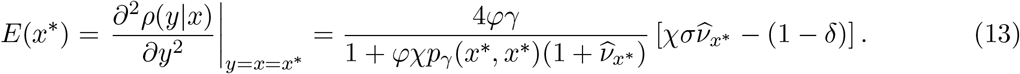

Specifically, *M*(*x*\*) < 0 corresponds to a singular point *x*\* being convergent stable, and *E*(*x*\*) < 0 corresponds to a singular point *x*\* being evolutionary stable, see also Supplementary Information S4

#### 2.2.2 Computer simulations

##### Stochasticity in adaptive dynamics

To assess the robustness of our analytical results to violation of some of the adaptive dynamics assumptions, we set up stochastic computer simulations to study the evolutionary dynamics of a population of initially monomorphic enantiostyly with phenotypic trait *x*. More specifically, in our simulations we included demographic stochasticity of emerging mutants, stochasticity in the selection and mutation process, and we relaxed the assumption of clear separation between population dynamic and evolutionary time scales. In order to do so, we used a computational algorithm developed by Saltini et al. (2023) and an adapted version of the code, see Supplementary Information S2 for details.

##### Population genetics

In our model we did not consider any explicit genetics nor did we allow for intermediate phenotypes resulting from cross-breeding. Therefore, we used population genetics simulations to assess whether evolutionary branching could occur in the case in which intermediate phenotypes can be generated by cross-breeding between plants with different phenotypic traits, when trait inheritance occurs through both male and female function. In these, plants have a Mendelian *E/e* locus which determines which of two sets of small effects loci (*x*_*E*_ or *x*_*e*_) is expressed. Each set itself is modelled as a single number here called *“meta-allele”*. See Supplementary Information S3 for details.

## 3 Results

### 3.1 Evolutionary dynamics

#### 3.1.1 Available evolutionary scenarios around x* = 1/2

Based on the signs of *M*(*x*\*) and *E*(*x*\*), we study the possible fates of a monomorphic population by investigating *x*\* = 1/2. If we combine Equations (12) for *M*(*x*\*) and (13) for *E*(*x*\*), we obtain

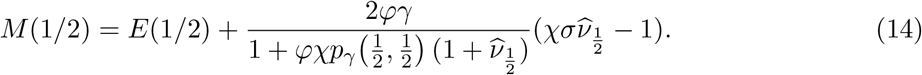

This implies that, if *E*(1/2) < 0, necessarily 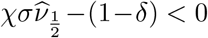 and, since 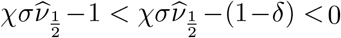, from Equation (12) it follows *E*(1/2) < 0 ⇒ *M*(1/2) < 0. This means that in our model, evolutionary stability also implies convergence stability (Figure 2A). Consequently, out of the four possible characterizations of the singular point *x*\* = 1/2 (Geritz et al., 1998), only three are available in our case (Figure 2A-C). If *M*(1/2) < 0 and *E*(1/2) < 0, *x*\* = 1/2 is at a maximum of the invasion fitness and, therefore, uninvadable by any local mutant. This is an end-point of the evolutionary dynamics and evolution results with a monomorphic population with 1:1 ratio of L- and R-flowers within individuals as the evolutionary stable strategy (ESS), see Figure 2A. From Equation (13), we note that such a condition is more easily met for low values of *δ*, i.e., for low fitness cost due to geitonogamy. The light grey area in Figure 2D is the area of coexistence and every pair (*x*_1_, *x*_2_) in the area of coexistence is a protected dimorphism. If a protected stylar dimorphism with two phenotypes in the area of coexistence occurs due to, for example, mutations of large phenotypic effect or immigration, such a dimorphism would be transient, as the coevolutionary dynamics would make the two resident trait values converge to *x* = 1/2, i.e. the monomorphic state (see Figure S3B).

If *E*(1/2) > 0 and *M*(1/2) < 0, then *x*\* = 1/2 is convergently stable but not evolutionarily stable (Figure 2B). This means that first directional selection drives the evolving trait *x* towards *x*\* = 1/2. Then, a monomorphic population splits, resulting in the emergence of a protected dimorphism, with the two resulting phenotypic traits undergoing disruptive selection and subsequent evolution leading to a dimorphic population toward *x* = 0 and *x* = 1 (Figure 2E). In this case, *x*\* = 1/2 is an evolutionary branching point. Note that, in our model, the invasion of a mutant with uniform handedness (either *y* = 0 or *y* = 1, corresponding to a large mutation step *ε* = ±1/2) into a population at the branching point *x*\* = 1/2 also leads to dimorphism (see Figures 2B,C and S3A). This demonstrates that our findings are robust under the relaxation of the assumption that mutations have small phenotypic effects.

Finally, if both *E*(1/2) > 0 and *M*(1/2) > 0, the strategy *x*\* = 1/2 is both convergent and evolutionary unstable (Figure 2C). In this case, *x*\* = 1/2 is a repeller of the evolutionary dynamics. With mutations of small phenotypic effect, the resident phenotype *x* evolves away from the singular point, resulting in a monomorphic population with trait either *x* = 0 or *x* = 1 (depending on the initial trait value), see Figure 2C. However, such a population is invadable from another part of the range, where the alternative strategy (*x* = 1 or *x* = 0, respectively) was present (Figure S4A). Furthermore, in the repeller regime, mutations of large phenotypic effect can result in the creation of stylar dimorphism, see Figure 2F and Figure S4B.

Note that complete inbreeding depression (*δ* = 1) implies evolutionary instability (*E*(1/2) > 0), suggesting that high levels of inbreeding depression resulting from geitonogamous selfing is a key factor in driving the evolution of dimorphic enantiostyly.

#### 3.1.2 Stochasticity in the demography and mutation-selection process

Figure 2G-I illustrates evolutionary outcomes for the simulated evolution of an initially monomorphic population for three different sets of model parameters. In Figure 2G, the monomorphic population evolves towards *x* = 1/2, where it then remains at a fitness peak. There, nearby mutants have a negative invasion fitness and, therefore, trait value *x* = 1/2 is an ESS.

In Figure 2H, after a short period of directional selection until *x* = 1/2, the monomorphic population undergoes evolutionary branching, resulting in the emergence of stylar dimorphism. Given the proximity of the two phenotypic traits emerging from evolutionary branching to the branching point *x* = 1/2, the lack of separation of time scales, and stochasticity in the mutation and selection process, subsequent evolutionary branching events occur resulting in a transient polymorphism with several coexisting phenotypic trait values. This diversity disappears when two branches evolve sufficiently far away from the branching point *x* = 1/2. Then, through negative frequency-dependent selection, evolution results in character displacement and eventually an evolutionary stable coalition, with the population exhibiting the two traits *x*_1_ = 0 and *x*_2_ = 1. However, if environmental changes occur such that the geometrical shape of the area of coexistence plot changes to something similar to Figure 2E, the evolution of the traits *x*_1_ and *x*_2_ resume. In such a case, the evolutionary trajectories of the two traits will converge to *x* = 1/2 and monomorphism is restored (Figure S3B).

In Figure 2I an initially monomorphic enantiostylous population evolves away from the singular point, resulting in a monomorphic population with fixed and uniform stylar orientation among all individuals (*x* = 0 or *x* = 1). This occurs because *x*\* = 1/2 is not convergent stable (*M*(1/2) > 0) and, in such a case, a monomorphic population evolves away from the singular point.

#### 3.1.3 Individual-based simulations including genetic system

To provide proof-of-principle that evolutionary branching through gradual evolution could originate in a population of sexually reproducing individuals, we performed population genetics simulations with the following genetic architecture: 1) individuals contain two meta-alleles, dubbed *x*_*e*_ and *x*_*E*_, corresponding to two sets of small effect loci, and a single diploid *E/e* locus, the latter similar to *Heteranthera missouriensis* (Jesson and Barrett, 2002*a*). *EE* and *Ee* individuals produce flowers according to their *x*_*E*_ meta-allele, *ee* individuals express *x*_*e*_. Figure 3A shows the outcome of a population genetic simulation with *f* = 12 flowers per plant in which an initially monomorphic population undergoes evolutionary branching resulting in the emergence of stylar dimorphism. In the first stage, where *x*_*e*_ and *x*_*E*_ were still similar, *f*_*E*_ - the allele frequency of *E*, fluctuated considerably. Later, when *x*_*e*_ and *x*_*E*_ had evolved sufficiently far away from one another, *f*_*E*_ stabilized around 0.3, i.e., corresponding to similar fractions of low-*x* and high-*x* individuals in the population. The great reduction in the fluctuation in *f*_*E*_ suggests that the *E/e* system is maintained by negative frequency-dependent selection when the two meta-alleles are sufficiently different. Genetic models considering many tightly linked loci (e.g., Bolnick and Doebeli, 2003) would likely yield very similar results (de Jong and Geritz, 2001).

**Figure 3:**
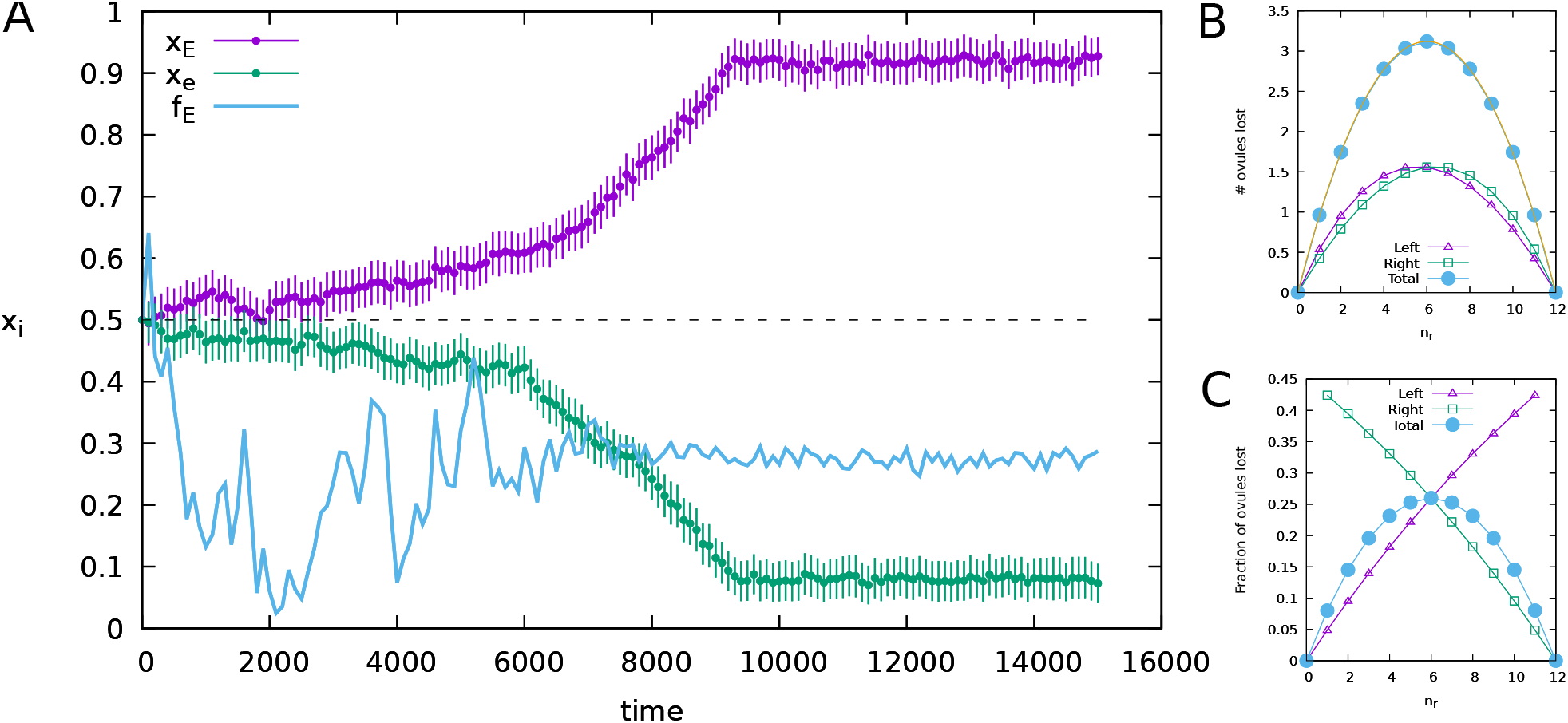
Evolutionary branching of an initially monomorphic population. (A) Outcome of a population genetics simulation where an initially monomorphic heterozygous population splits into a dimorphic population. Green: average *x*-value of the *x*_*e*_-metagene ± standard deviation, purple: average similar for *x*_*E*_-metagene. Blue line: population allele frequency *f*_*E*_ of the *E* -allele. Note that the fluctuations in *f*_*E*_ decrease after *x*_*e*_ and *x*_*E*_ are sufficiently diverged, indicating that from then on, the *E/e*-system is maintained by negative frequency dependent selection, resulting in roughly equal fractions of *x*_*e*_ and *x*_*E*_ expressing individuals. (B) Expected number of ovules lost due to geitonogamy with the parameters used in the simulations, split for L- and R-flowers and summed. Yellow curve is a parabola calculated to go to all points of the total. (C) Relative ovule loss corresponding to B. Parameters: *f* = 12, *v* = 12, *ζ* = 2, *λ* = 0.2, *ε* = 0.2.

Our simulations show that branching is possible by gradual evolution but, due to stochasticity, in a fraction of simulations in the branching regime, either the *E* or *e* allele was lost during the initial stage of a simulation, after which selection drove *x* of the remaining allele to around 0.5.

### 3.2 Bifurcation analysis

To assess how characterization of the singular point *x*\* = 1/2 (and, therefore, the evolutionary outcome) depends on numerical values of the model parameters, we performed bifurcation analyses at *x* = 1/2. This was conducted by changing, independently of each other, the cost of geitonogamy *δ*, and the reduction in pollination efficiency for pollen transferred between flowers of the same stylar orientation *γ* for multiple values of the reproductive rate *φ* (Fig. 4), the competition coefficient *σ* (Fig. 5), and the handling time *χ* (Fig. S1).

**Figure 4:**
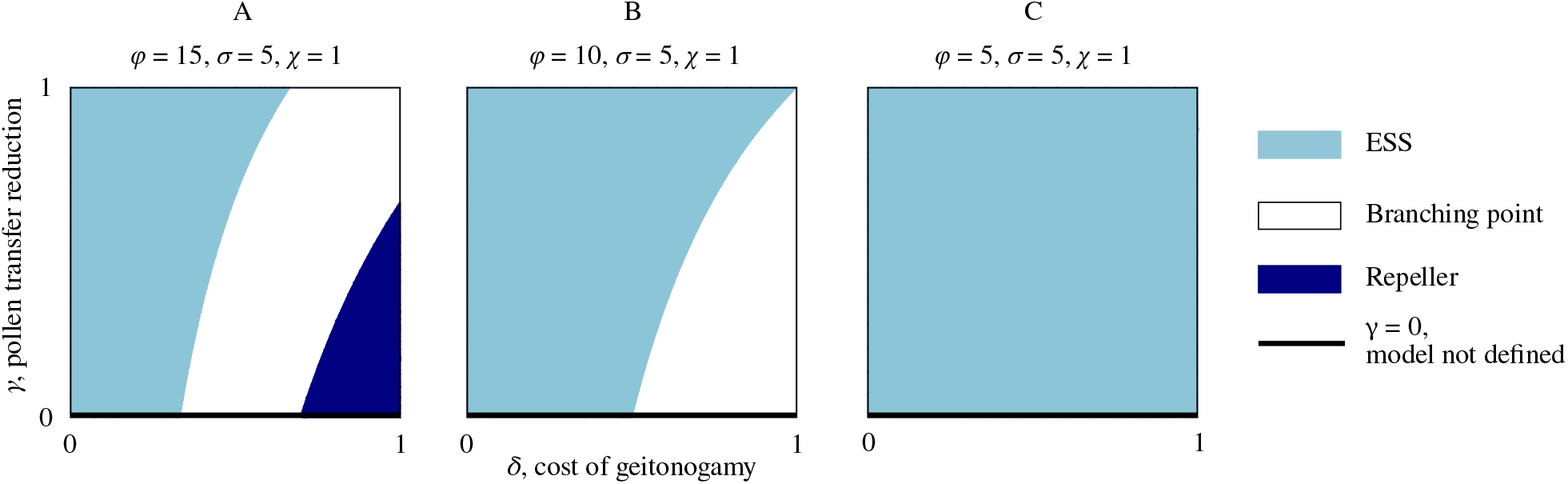
High reproductive rate favours the evolution of dimorphic enantiostyly. Characterization of the singular point *x*\* = 1/2 with three bifurcation diagrams in the *δ* and *γ* parameters for decreasing (left to right) values of the reproductive rate *φ*. (A) Depending on the combination of values of *δ* and *γ*, for high reproductive rate the evolutionary outcome of the model can be an ESS for the monomorphic population (light blue), a branching point with consequent emergence of dimorphism (white), and a repeller of the evolutionary dynamics (dark blue) in which, for most initial conditions, mutations of small effect result in a monomorphic population with uniform direction for the stylar deflection among individuals, i.e., with trait value either *x* = 0 or *x* = 1. (B) By decreasing the reproductive rate, the parameter regime that leads to an evolutionary repeller shifts beyond *γ* > 1, leaving only an ESS or branching point as possible characterizations of *x*\* = 1/2. (C) For lower value of the reproductive rate *φ*, the singular point *x*\* = 1/2 is an ESS for any biologically relevant combination of *δ* and *γ*.

**Figure 5:**
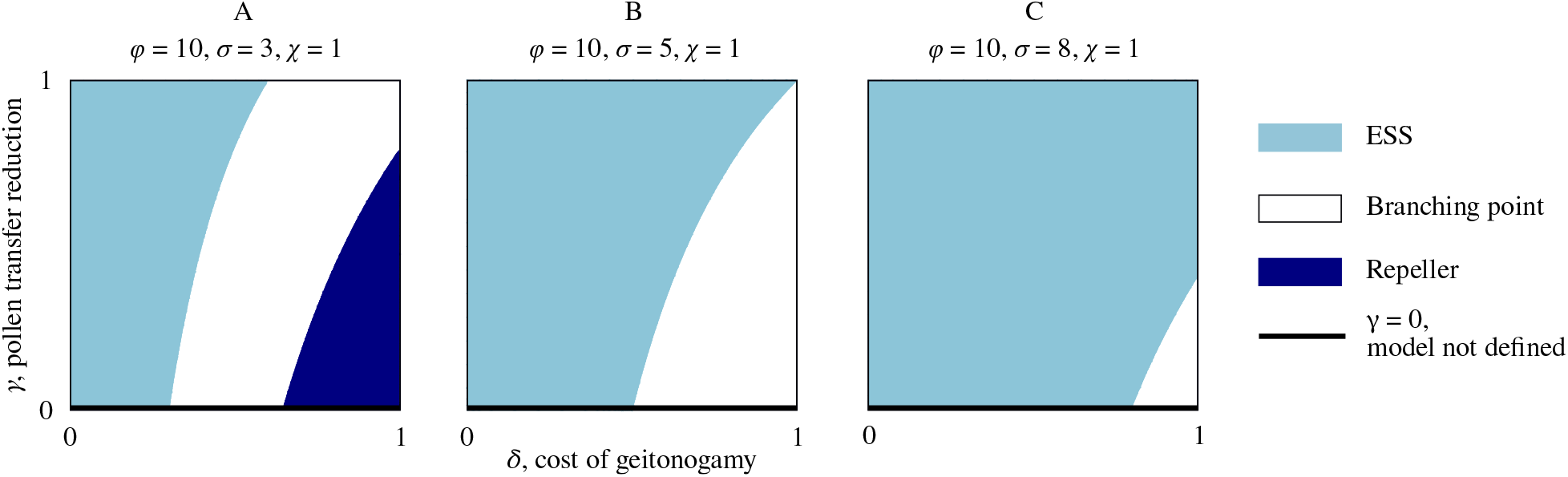
Increased competition reduces the scope for evolving dimorphic enantiostyly. Characterization of the singular point *x*\* = 1/2 with three bifurcation diagrams in the *δ* and *γ* parameters for increasing (left to right) values of the competition coefficient *σ* for shared abiotic resources. (A) Depending on the combination of values of *δ* and *γ*, for weak competition three evolutionary outcomes are possible: an ESS for the monomorphic population (light blue), a branching point with consequent emergence of dimorphism (white), and a repeller of the evolutionary dynamics (dark blue) in which, for most initial conditions, mutations of small effect result in a monomorphic population with uniform direction for stylar deflection among individuals, i.e., with trait value either x = 0 or x = 1. (B) By increasing the competition coefficient, the parameter regime that leads to an evolutionary repeller shifts beyond *γ* > 1, leaving only ESS or branching point as possible characterization of *x*\* = 1/2. (C) For higher value of the competition coefficient *σ*, the range of values for *δ* and *γ* such that *x*\* = 1/2 is a branching point shrinks, and evolutionary branching can only occur for very high *δ* and very low *γ*, whereas for other combination of *δ* and *γ* the singular point *x*\* = 1/2 is an ESS.

Figure 4 illustrates three bifurcation diagrams for the parameters *δ* and *γ* for decreasing values of the reproductive rate *φ*, calculated at the singular point *x*\* = 1/2. In particular, Figure 4A shows that a gradual increase of *δ* with a fixed, low value of *γ* changes *x*\* = 1/2 from an ESS to a branching point and, for a very high geitonogamy cost, to a repeller of the evolutionary dynamics. Note that the region in Figure 4A where *x*\* = 1/2 is not evolutionary stable (branching point and repeller) is broader than the same region both in Figure 4B, where the singular point cannot be a repeller, as the repeller regime has shifted beyond *γ* > 1, and in Figure 4C, where any combination of *δ* and *γ* corresponds to an ESS for the singular point. Therefore, the broader scope for evolutionary instability and a branching point is associated with higher values of the reproductive rate *φ*. Note that *φ* increases with increasing daily display size *f*, more efficient pollen-to-seed conversion *c*, and/or increasing pollinator activity *r*_*p*_. In this case, a monomorphic population can undergo evolutionary branching into two phenotypes with fixed, reciprocal handedness, or evolve towards a uniform direction for stylar deflection among individuals (bottom right corner of Figure 4A). In Figure 4A, whereas lower values of *γ* are linked to a broader range of values for *δ* such that *x*\* = 1/2 is evolutionary unstable, the scope for evolutionary branching becomes slightly narrower compared to higher values of *γ* because of the appearance of the repeller regime. In contrast, a low *φ* value is associated with an increased area in the (*δ, γ*) plane where the strategy *x*\* = 1/2 is evolutionary stable. In this case, a monomorphic population with a 1:1 proportion of L- and R-flowers within individuals is the ESS.

Similarly, strong competition for shared abiotic resources *σ* leads to a widening of the region of the (*δ, γ*) space where the singular point *x*\* = 1/2 is evolutionary stable, whereas weak competition results in conditions in which evolutionary branching and an evolutionary repeller state are more likely to occur (Figure 5).

Finally, Figure S1 illustrates that changing the numerical value of the handling time *χ* does not change the characterization of the singular point. Thus, while the numerical value of both *M*(1/2) and *E*(1/2) changes significantly in response to changes of *χ* and, consequently, so does the speed of gradual evolution, the sign of the two functions is fully independent of the changes in handling time.

## 4 Discussion

The primary goal of our study was to investigate phenotypic diversification of a plant population with monomorphic enantiostyly, in which all individuals have similar mixed stylar conditions of L- and R-flowers, into a dimorphic population in which individual plants are fixed for either L- or R-flowers. This evolutionary transition in sexual system involves the origin of a true genetic polymorphism in which individuals represent phenotypically distinct floral morphs maintained in populations by negative frequency-dependent selection between L- and R-plants. We approached this problem by employing the modelling framework of adaptive dynamics to study how the ecological conditions influencing the dynamics of a monomorphic population could cause the emergence of stylar dimorphism under the assumption of gradual adaptive evolution. Nevertheless, our results remain robust even when allowing for abrupt evolutionary changes, i.e., they hold if we assume mutations with large phenotypic effects. We showed, using population genetic simulations of a similar model, that this transition is possible assuming a plausible genetic system underlying the relevant trait. Such simulations also revealed that stochasticity often led to the loss of either the *E* or *e* allele, impeding the shift from monomorphism to dimorphism despite otherwise favourable conditions. This may be one of the explanations why dimorphic enantiostyly is rare.

Our investigation of the evolution of dimorphic enantiostyly differs from most earlier theoretical models of the evolution of plant sexual polymorphisms, including dioecy and heterostyly. Theory on this topic has most often employed an explicit population genetic framework (Charlesworth and Charlesworth, 1979*a*; Lewis, 1941) and/or has used phenotypic selection models based on sex allocation theory and fitness gain curves (Charnov, 1982; Lloyd, 1982). In common with our study, these models have generally concerned transitions from monomorphic to dimorphic conditions; but because of intrinsic differences in the floral biology of these contrasting sexual systems, the models vary in the specific parameters investigated. Dioecy commonly evolves from monoecy (Pannell and Jordan, 2022; Renner and Ricklefs, 1995) and some models of this transition have been based on quantitative changes in male or female fertility through gradual alteration in the relative proportions of female and male flowers in ancestral monoecious populations (Charlesworth and Charlesworth, 1978). Key parameters driving selection of unisexual individuals usually involve the selfing rate and Inbreeding depression of hermaphrodite offspring (Charlesworth, 1999). To our knowledge, the only theory that has explicitly used the adaptive dynamics approach for sexual systems transitions investigated the role of geitonogamy in the gradual evolution of dioecy (de Jong and Geritz, 2001). Significantly, these authors demonstrated that the intensity of inbreeding depression required for the evolution of dioecy differed depending on whether the plants considered were animal versus wind-pollinated. Because all flowers in monoecious and dioecious sexual systems are unisexual rather than hermaphroditic, geitonogamous pollen transfer is much simplified compared to our case where we investigated the consequences of pollen transfer both between and within L- and R-handed flowers.

The evolution of the stylar polymorphism distyly also involves ancestral monomorphic populations initially fixed for a single stylar condition. In this case the establishment of stylar dimorphism results from the spread of variants with modified sex-organ arrangement within flowers. The two most widely accepted models for the evolution of distyly differ in the relative emphasis that they place on inbreeding depression (Charlesworth and Charlesworth, 1979*b*) versus the advantages of disassortative pollen transfer (Lloyd and Webb, 1992*b*). However, both models differ fundamentally from our approach because in heterostylous species, and all other stylar polymorphisms with the exception of enantiostyly, individuals in ancestral monomorphic populations do not produce the alternate stylar conditions. In this respect the transition we have investigated shares more similarities with the monoecy – dioecy transition since in both cases the developmental machinery required to produce the two flower types (e.g. female and male or L- and R-styled) already occurs in ancestral populations.

Note that our study specifically focused on the evolution of dimorphic enantiostyly in relation to the L:R ratio of flower handedness, assuming an already fixed and evolutionary stable degree of stigma-anther separation (herkogamy). Nevertheless, we believe that the co-evolution of herkogamy and the ratio of L:R flowers would be worthy of further investigation.

### 4.1 Reductions in geitonogamy can promote transitions to dimorphism

We found that facilitative interactions between individuals with different stylar phenotypes, mediated by the pollinating function *p*_*γ*_, resulted in the evolution of dimorphism from monomorphism through disruptive selection on the proportion of L- and R-flowers within a plant (Fig. 2H). This transition occurred when the fitness costs of geitonogamy owing to inbreeding depression were high enough to favour phenotypes fixed for stylar condition (Figs. 4, 5). Experimental studies of pollen transfer have demonstrated that sufficiently large bees can promote pollinations between flowers of opposite stylar orientation (Minnaar and Anderson, 2021). As a result, levels of geitonogamous selfing should be reduced significantly for dimorphically enantiostylous plants in comparison with those in monomorphic populations as the latter have both L- and R-flowers and are thus prone to between-flower self-pollination (Jesson and Barrett, 2002*b*). In agreement with our results, previous studies have also proposed that the costs of geitonogamy can play an important role in floral evolution, including the evolution of sexual polymorphisms in reproductive traits (de Jong and Geritz, 2001; Harder and Barrett, 1995; Jesson and Barrett, 2005). Geitonogamy may be a ubiquitous outcome in species in which daily display sizes involve numerous flowers. It is therefore unsurprising, given the potential female (inbreeding depression) and male (pollen discounting) mating costs of geitonogamy, that animal-pollinated plant populations have evolved diverse floral mechanisms such as enantiostyly to limit pollen transfer between flowers within a plant.

### 4.2 Ecological factors influence invasion scenarios and evolutionary branching

Our model predicted that a high reproductive rate which, in turn, depended on the size of the daily floral display, the pollen-to-seed conversion efficiency, and pollinator activity, broadened the scope for the emergence of evolutionary branching in the evolving trait (stylar orientation) we considered (Fig. 4). Conversely, strong competition for shared resources such as soil moisture or nutrients impeded evolutionary branching and instead favoured the evolutionary stability of monomorphic enantiostyly (Fig. 5). Unfortunately, explicit comparative analyses of traits and ecological characteristics between the two distinct forms of mirror-image flowers may not be especially revealing because of the limited number of evolutionary transitions to the dimorphic state in angiosperms.

In their theoretical investigation of the evolution of enantiostyly, Jesson et al. (2003*a*) proposed that dimorphism evolved from monomorphism through two separate and sequential invasions of variants with fixed, reciprocal handedness. By making this assumption they implicitly assumed that after the first invasion (assuming that they were not simultaneous, which seems improbable), the resident monomorphic population with mixed handedness was not replaced by the variant. If this did occur after the replacement of the resident, the persistence of the variant would be contingent on the efficiency of pollen transfer between flowers with the same stylar orientation, a scenario not discussed by Jesson et al. (2003*a*). Assuming that coexistence between a population of plants with mixed handedness and a variant with fixed handedness was possible, then our model predicts that because of negative frequency-dependent selection imposed by the reproductive benefits of between-flower type pollination, the population of plants with mixed stylar orientation would gradually evolve to dimorphic enantiostyly as the proportion of L- or R-flowers on mixed plants would likely change over time to reciprocally match the variant (Fig. 2E, S3A). In summary, our model supports the Jesson et al. (2003*a*) scenario of two sequential invasions with fixed, reciprocal handedness (Fig. 2B and E), and further sheds light on the evolutionary forces in between their hypothesised sequential invasions.

Significantly, our models revealed that inbreeding depression following geitonogamy combined with the presence of pollinators that can efficiently mediate pollen transfer among flowers of both stylar orientations (low *γ* values) broadened the scope for evolutionary instability of an initially monomorphic population. However, the range of values for the fitness cost due to geitonogamy resulting in evolutionary branching was slightly decreased. This situation provided an advantage to the evolutionary repeller resulting in a monomorphic population with the direction of stylar deflection uniform among all individuals (Figs. 4, 5). Although relatively uncommon, this type of herkogamous stylar condition occurs in several angiosperm families (e.g., Gentianaceae). Also five populations of *Tenicroa*(= *Drimia*) *exuviate* of the Hyacinthaceae (Jesson and Barrett, 2003) have been reported with exclusively right-deflected styles. This type of herkogamy can result in lower pollen transfer between flowers and, hence, in reduced probability of geitonogamous selfing.

The viability of this evolutionary strategy depends on whether the ecological and environmental conditions allow for a sufficient rate of pollination. Indeed, the likelihood of the evolution of populations with a single stylar orientation may be greater in situations where there is weak competition for abiotic resources between individuals as this may positively increase plant density and promote and increased outcrossing. Thus, our models suggest that the existence of the stylar deflection of the type that occurs in *Tenicroa* might be attained through selection for a single stylar orientation rather than through the breakdown of dimorphic enantiostyly and loss of either the L- or R-phenotype from populations.

Finally, our model predicted that starting from a monomorphic state, complete inbreeding depression (*δ* = 1) leads to evolutionary branching and the origin of dimorphic enantiostyly or, alternatively, evolution towards a population fixed for a single stylar orientation. For this to occur, however, the population needs to exhibit a large enough floral display to cause geitonogamy, and/or there is weak competition among plants for resources. This is because the viability of a non-zero equilibrium density when fitness costs due to geitonogamy are maximum depend on whether individuals still have significant mating opportunities, i.e., the density of plants is sufficiently high to allow for adequate transfer of pollen(see Supplementary Information S1).

### 4.3 High reproductive rate and weak competition favour the emergence of dimorphism

Our bifurcation analysis indicated that a high reproductive rate and/or a weak competition coefficient broadened the scope for evolutionary branching (Figs. 4 and 5). In our model, the reproductive rate was influenced by several factors. Specifically, the daily display size had a quadratic effect whereas both the pollen-to-seed conversion coefficient and the pollinator visitation rate had a linear effect on the reproductive rate. Therefore, the absence of dimorphic enantiostyly in some lineages with monomorphic enantiostyly may be attributed to selective constraints associated with small daily floral display size, as suggested by Barrett et al. (2000). Thus, for dimorphism to evolve, a monomorphic population may first have to develop larger floral displays.

A recent study found a correlation between inbreeding depression and decreased floral longevity (Spigler and Charles, 2023). If decreased floral longevity is not counterbalanced by an increased rate of flowering, it would likely result in decreased daily floral display size. Such a daily floral display size, in our models, is positively correlated to the emergence of dimorphic enantiostyly. In our models the key driver of evolutionary branching is geitonogamy, thus the observation that inbreeding depression may limit the size of daily floral displays suggests the existence of additional constraints on the emergence of dimorphism. Our models also suggest that the quantity and quality of pollinators visiting populations may also prevent the emergence of dimorphism from monomorphism by decreasing reproductive rate. On the one hand, pollinators capable of efficiently transferring pollen between flowers, regardless of the direction of stylar deflection, broaden the scope for the evolutionary instability of monomorphism. Alternatively, they may actually weaken selection for optimal L:R ratios and prevent a population from evolving towards dimorphism (Fig. S2).

It is also possible that limited pollinator service could potentially trigger selection of plants with larger daily floral displays (Sletvold et al., 2010), as large displays generally attract more pollinators (Grindeland et al., 2005). Also, as occurs in the orchid *Satyrium longicauda*, there is evidence of plasticity in display size in response to variation in pollinator service (Harder and Johnson, 2005; Spigler, 2017). In a future study, it would be interesting to explore whether changes in daily display size (due to either plasticity or adaptive evolution) could affect the emergence of dimorphism in mirror-image flowers.

Both high reproductive rate and weak competition for shared resources positively influenced the density of individuals, with the competition coefficient ultimately being the most influential parameter in controlling density. This relationship means that plant density arises naturally from population dynamics considerations, rather than being included in the model as an independent parameter. A high density of individuals provides more mating opportunities compared to a situation in which individuals are sparsely distributed. Indeed, the latter case would be more likely to favour increased self-fertilization (Karron et al., 1995; Treuren et al., 1993). Thus, the demographic features of populations may be crucial for dimorphic enantiostyly to evolve, as low-density conditions are likely to result in a decreased probability of outcrossing and a reduction in the number of potential mating partners of the alternate stylar morph compared with a monomorphic population in which all individuals can potentially mate with one another. In this sense, a sparse population may not provide sufficient opportunities for outcrossing, making dimorphic enantiostyly a disadvantageous sexual system in terms of reproductive fitness.

Our model predicted that under restricted circumstances reversions from dimorphic enantiostyly to monomorphic enantiostyly might occur. Changes in the ecology of populations, such as a decline in pollinator service or increased competition for shared resources, were found to decrease the reproductive rate or increase the competition coefficient. This had the potential to bring a dimorphic population closer to the monomorphic regime, or even fully into it, with evolutionary trajectories that converge to *x* = 1/2 (Fig. 2D and S3B).

### 4.4 Enantiostyly as a case study for evolutionary branching driven by ecological facilitation

Whether evolution in our models results in a monomorphic or a dimorphic enantiostylous population strongly depends on the pollination function. This dependency occurs because this parameter facilitates reproductive connections between flowers within and among individuals through mating. In most adaptive dynamic models, inter- and intra-specific interactions are modelled through trait-dependent competition coefficients, representing competition for shared resources. In these circumstances, negative frequency-dependent selection serves to decrease intra-specific competition through evolutionary branching, with disruptive selection and subsequent character displacement minimizing inter-specific competition and ultimately maximizing fitness for specific trait patterns. In contrast to this commonly studied scenario, we used a facilitative interaction function, i.e., the pollination function. In this context, the maximization of fitness is achieved through the maximization of facilitation, rather than the minimization of competition (Fig. S2A,C). This condition is particularly true when specialized pollinators visit enantiostylous flowers, i.e., pollinators that are far more effective in dispersing pollen between flowers of the opposite rather than the same stylar orientation (high *γ* values, Fig. S2B). The resulting maximization of positive interaction leads to evolutionary branching, followed by disruptive selection with character displacement and the origin of dimorphism.

The idea that different types of ecological interactions other than competition, such as facilitation, cooperation, or mutualism can drive evolutionary diversification by generating resources and therefore niches is supported by both empirical (Blake et al., 2021; Turner et al., 1996) and theoretical (de Jong and Geritz, 2001; Doebeli, 2002; Doebeli and Dieckmann, 2000; Wechsler and Bascompte, 2022) evidence. However, the underlying mechanisms promoting evolutionary diversification through interactions that are cooperative or facilitative remain relatively poorly understood (Bird et al., 2019; Day and Young, 2004; Hardy et al., 2020), despite the widespread occurrence of these types of interaction in diverse ecosystems including plant-insect, bacterial, and microbial biofilm networks (Blake et al., 2021; Poltak and Cooper, 2010). Our model is a striking example of evolutionary diversification resulting from positive ecological interactions and may serve to stimulate future investigations of the interplay between facilitative interactions and the potential for evolutionary branching.

## Acknowledgement

This work was supported by the Human Frontier Science Program (HFSP), grant no. RGP0036/2021.

## Author contribution

MS and EED designed the models with input from SCHB, and carried out the formal analysis. All authors conceived the study, interpreted the results, and wrote and edited the manuscript.

## Supplementary Information

### S1 Stationary solution of the population dynamics of a monomorphic population

We calculate the size of the population at the dynamic equilibrium 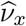 by solving for d*ν*_*x*_/d*τ* = 0 from Equation (7). The system has two biologically relevant solutions: the trivial solution 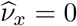 and

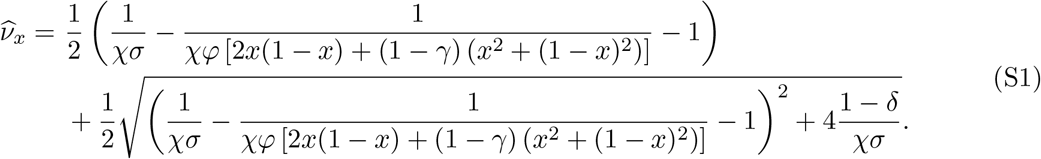

A third solution of the equation d*ν*_*x*_/d*τ* = 0 exists, with 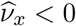 which is not realistic and, therefore, it is rejected. Isocline analysis reveals that, for *x* ∈ (0, 1), the solution 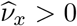 is an attractor of the population dynamics, whereas 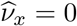 is a repeller. This means that, for any *ν*_*x*_ at time *τ* = 0, the population dynamics always converges to 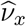.

Note that, if *γ* = 1, Equation (S1) has two diverging terms if *x* = 0 or *x* = 1. By taking the limit for *γ* → 1, we can show that in this case 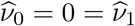. Intuitively, this means that a monomorphic population of plants with a single, fixed handedness, can only survive if pollinators can also transfer pollen between flowers with of same stylar orientation.

Similarly, for *δ* = 1, the term outside the squared root of Equation (S1) must be positive, to obtain a positive equilibrium density. This condition is more likely met for lower values of *σ* (weak competition) and high values of *φ* (high reproductive rate). This shows that, if self-fertilization is particularly expensive in terms of fitness loss, a population can only survive if the competition for shared abiotic resources is weak and/or the reproductive rate is high.

### S2 Description of the computational algorithm

To study the robustness of our analytical results to violation of assumption inherent to the adaptive dynamics framework, we set up stochastic computer simulations where we include demographic stochasticity in the population dynamics of new variants, and we relax the assumption of strict separation between the fast ecological and the slow evolutionary time scale. The inclusion of demographic stochasticity for newly-occurred mutants, when their density is still low compared to the density of resident phenotypes, implies that even beneficial mutations have a finite probability to go extinct and not establish in the population. As a consequence, if more phenotypes coexist in a given population, mutation and selection is a process subject to stochasticity as the establishment of beneficial mutations depends on both the probability that a new mutant occurs from one of the phenotypes of the population, and on the probability that such mutant survives demographic stochasticity when it still is rare (Dieckmann & Law, 1996).

More specifically, suppose that a mutant with trait *y*_*i*_ occurs from the resident *x*_*i*_ in a population consisting of *n* phenotypes with traits *x*_1_, …, *x*_*n*_. Then, the rate Ω at which *y*_*i*_ invades the resident population is

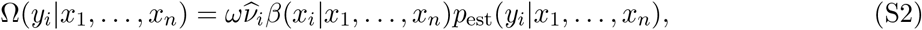

(Dieckmann & Law, 1996; Champagnat et al., 2006). In the last equation, *ω* is the *per-capita* probability to produce a mutation, 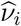 is the equilibrium density of the phenotype from which the mutant has occurred, *β* is the *per-capita* reproductive rate of the phenotype *i* and is defined in Equation (9), and

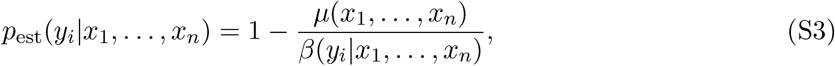

is the establishment probability of the mutant phenotype with trait *y*_*i*_, i.e., the probability that the mutant survives demographic stochasticity and successfully invades the population (Allen, 2010; Saltini et al., 2023), and where *µ* is the *per-capita* mortality rate defined in Equation (10).

From Equation (S2) it follows that the rate at which new mutations establish depends on the phenotypic traits in the populations which are variable quantities subject to evolutionary dynamics. Therefore, Ω is not a fixed rate and, consequently, the time that passes between different consecutive successful mutations is a stochastic quantity. Here, as we are not interested in any time-dependent analysis but only in the conditions that lead to different possible evolutionary outcomes, we use an event-driven approach and we measure time in unit of successful mutations.

The simulation algorithm, developed by Saltini et al. (2023) is programmed in C++ and works as follows:

1. Simulations are initiated with a monomorphic population with trait *x* = 0.25.
2. Numerically solve Equation (7) with Euler’s forward method with a time increment of *τ* = 0.01 until either the difference between the density of every phenotype before and after an iteration of the population dynamics is smaller than 10^−9^ or after a maximum of 10^5^ iterations has been reached. Note that, by imposing a cutoff to the population dynamics after 10^5^ iterations, we violate the strict separation between the fast ecological and the slow evolutionary time scale.
3. Remove from the population any phenotypes with density lower than 10^−3^.
4. For each of the *n* phenotypes *x*_*i*_ the population consists of, determine the two mutant traits *y*_*i*_ = *x*_*i*_ ± *ϵ*, with *ϵ* = 0.01.
5. Assign to each possible mutant the probability Ω(*y*_*i*_|*x*_1_, …, *x*_*n*_)/ Σ_{*y*}_ Ω(*y*|*x*_1_, …, *x*_*n*_) to be the next mutant to successfully invade the population. The summation in {*y*} runs over all 2*n* possible mutations that can occur in a population consisting of *n* phenotypes.
6. Based on these probabilities, stochastically select the next variant to be added to the population.
7. If the selected mutant has trait *y* > 1 or *y* < 0, reject the selection and return to (5).
8. Calculate the establishment probability for a mutant in the opposite direction compared to the selected mutant (i.e., if the selected mutant is *y*_*i*_ = *x*_*i*_ + *ε*, then calculate the establishment probability of *y*_*i*_ = *x*_*i*_ − *ε*).
  (8.a) If such establishment probability is also positive, add the selected mutant to the population and maintain the resident phenotypes. In this case, a lineage has split into two, and we refer to this event as evolutionary branching.
  (8.b) If such establishment probability is negative, replace the resident phenotype with trait *x*_*i*_ with the selected mutant with trait *y*_*i*_.
9. Reiterate the procedure starting from (2), until no new mutants can successfully invade.

### S3 Population genetics simulations

As trait *x* is both subject to evolution and shaping it, we performed population dynamic simulations to test whether the evolutionary trajectory of branching would be available in practice. For this, we developed python simulations of plants competing for a single resource (space in computer memory) and reproducing via explicit pollination.

The starting population consisted of *N* = 2000 plants. Each of the plants produced a total of *f* = 12 flowers, a fraction *x* of which right-handed, following a binomial distribution. From the initial number of flowers per plant, we assume that a fraction *g* is lost depending on the plant’s display. We first calculate the expected amount of geitonogamous pollen *g*_*L*_ and *g*_*R*_ received by individual L and R flowers during a single pollinator visit, respectively, based on the model by Jesson and Barrett (2005).

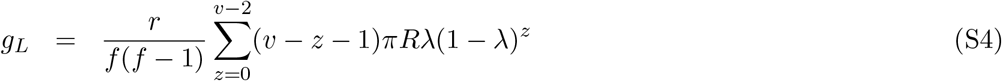

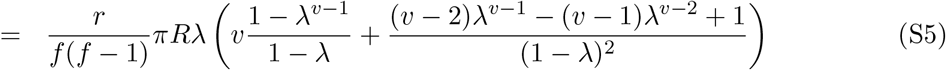

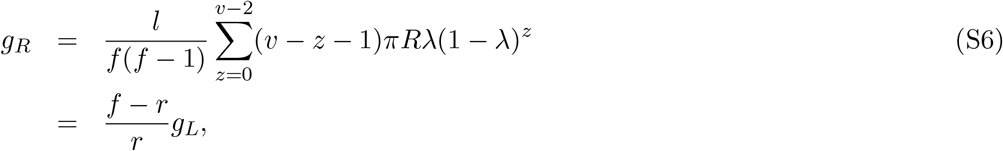

with *R* the number of pollen grains removed from each flower, *π* the fraction of those deposited on the pollinator’s body that are thus available for pollination, *λ* the fraction of pollen lost from the pollinator’s body at each flower contact “due to the vagaries of pollination”, *v* the number of flowers visited per pollinator visit (*v* ≤ *f*), *l* and *r* the number of left- and right-handed flowers on the plant, respectively. Note that, in the original model by Jesson and Barrett (2005), the fraction of pollen lost from the pollinator’s body is denoted by *ρ* which, in our adaptive dynamics model, denotes the invasion fitness.

For simplicity, we assume that a flower has one ovule. We assume that the probability of geitonogamy depending on self-pollen exposure is exponential with rate *q*, so the number of L-ovules (and R) fertilized by geitonogamy is Poisson distributed. These are lost with probability *δ* as in Equation 4.

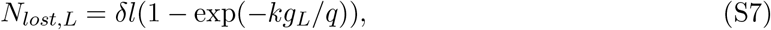

and similar for R, and *k* the number of pollinator visits. Note that parameters *πRk/q* ≡ *ζ* only occur together in this single combination, rendering them effectively a single parameter in this simulation. Moreover, the expected number of L+R ovules lost forms a perfect parabola, meaning the shape of the selective profile on *x* is the same as in the adaptive dynamics model, with tunable strength. Here, we use *ζ* = 2, *v* = 12, *λ* = 0.2.

With these parameters, a 6L-6R plant, for example, on average looses 1.56 L-ovules and 1.56 R-ovules, whereas a 11L-1R plant looses 0.54 and 0.42, respectively, and a 12L-0R none. Following Jesson and Barrett (2005), pollen loss is independent of flower orientation. We, therefore, keep lists of references to both potential male and female parents, with ovule loss only affecting the latter. All four lists are independently shuffled.

Given the total numbers of L and R ovules thus remaining, the number of L-L, Lmale-Rfemale, Rmale-Lfemale and R-R pollinations is calculated proportional to Equation 1, with *γ* = 0.7. The maximum number of attempted pollinations is *N*. A fraction *γ*(*f*_*LL*_ + *f*_*RR*_) of potential pollinations is not executed due to the inefficiency of intramorph pollination. The parents for the remaining crosses are then simply taken starting from the top of the respective lists, until the maximum of pollinations is reached. Upon reaching the bottom of a pollen list (either L or R)-list, the respective list is shuffled again, to avoid locking into the same pairs of crosses. Each ovule can produce at most one offspring.

Plants are diploid, with two *x*-metagenes, *x*_*E*_ and *x*_*e*_, taken as the result of many quantitative loci. The plant also has a single *E* -locus, reflecting the genetic system in *H. missouriensis* Barrett (2002*a*), with possible alleles *E* and *e. EE* and *Ee* plants express *x*_*E*_ as their *x*-trait, whereas *ee* plants express *x*_*e*_. Whether *x*_*E*_ or *x*_*e*_ will eventually code for right-handedness in a particular simulation, if evolutionary branching occurs, is purely coincidental. The value of the *x*-metagenes is inherited from parents 1 and 2 as *x*_*i*_ = *U*(max(0, min(*x*_*i*,1_, *x*_*i*,2_) − *ε*), min(1, max(*x*_*i*,1_, *x*_*i*,2_) + *ε*)), where *i* ∈ {*E, e*}, *U* represents the uniform distribution and *ε* = 0.02 is the maximum mutation step size. All plants are initialized *Ee* with *x*_*E*_ = *x*_*e*_ = 0.5.

At the start of each time step, a random 20% of the existing population dies. The remaining plants reproduce with a maximum of *N* new individuals. After that, population size is reduced to a maximum of *N* and a new time step starts, until *T* = 15000 or *T* = 20000.

### S4 Adaptive dynamics

We investigated gradual adaptive evolution of the phenotypic trait *x*. If a mutant invades and replaces a resident trait, it becomes the new resident and, in turn, can be invaded and replaced by a consecutive mutant (trait substitution sequence, see Figure 2G for an example).

#### The selection gradient

The direction and speed at which this process occurs is described by the selection gradient *s*(*x*), defined as

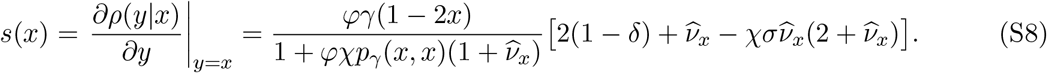

When the selection gradient is positive, the evolution of the trait *x* proceeds to increase its value, while a negative selection gradient leads to evolutionary changes in trait *x* that decrease its value.

#### Singular points

The singular points *x*\* of the evolutionary dynamics are trait values at which the selection gradient vanishes. These points correspond to resident phenotypes that are either at the maximum or at the minimum of the invasion fitness. Therefore, they satisfy the condition *s*(*x*\*) = 0. It can be verified that *x*\* = 1/2 is a singular point of the evolutionary dynamics, see Figure 2A-C. Note that *x*\* = 1/2 corresponds to a 1:1 ratio of L- and R-flowers, the typical condition found in populations of enantiostylous plant (Jesson and Barrett, 2003). In principle, our model permits other singular points, i.e., *x*\* values such that

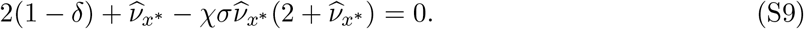

However, given our interest in study the emergence of polymorphism in enantiostyly starting from a 1:1 ratio of flowers with different handedness on a plan, we will only focus on the characterization of the trait *x*\* = 1/2.

#### Convergence stability

The first derivative of the selection gradient,

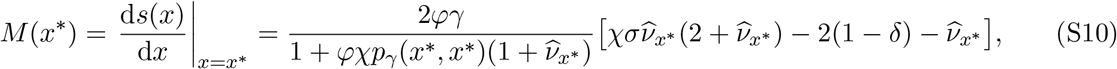

determines whether *x*\* is an attractor or a repeller of the evolutionary dynamics. Specifically, if *M*(*x*\*) < 0, the singular point *x*\* is convergently stable, and is an attractor of the evolutionary dynamics. In this case, directional selection proceeds towards *x*\*, see red arrows in Figure 2A,B. Conversely, *M*(*x*\*) > 0 implies that any trait substitution sequence for a resident phenotype *x* proceeds away from *x*\*, see red arrows in Figure 2C.

#### Evolutionary stability

The evolutionary stability of the singular point is determined by the sign of

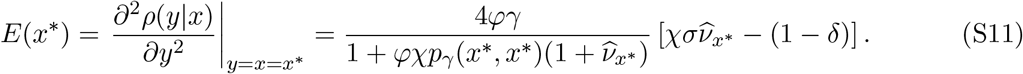

If *E*(*x*\*) < 0, then *x*\* is at a fitness maximum and is uninvadable by any nearby mutant and is, therefore, evolutionary stable. Instead, if *E*(*x*\*) > 0, then *x*\* is at a fitness minimum and nearby mutants from any direction can invade but with persistence of the resident trait *x*\* = 1/2 (Figure 2E,F); such a strategy is, therefore, evolutionary unstable.

### S5 Supplementary figures

**Figure S1:**
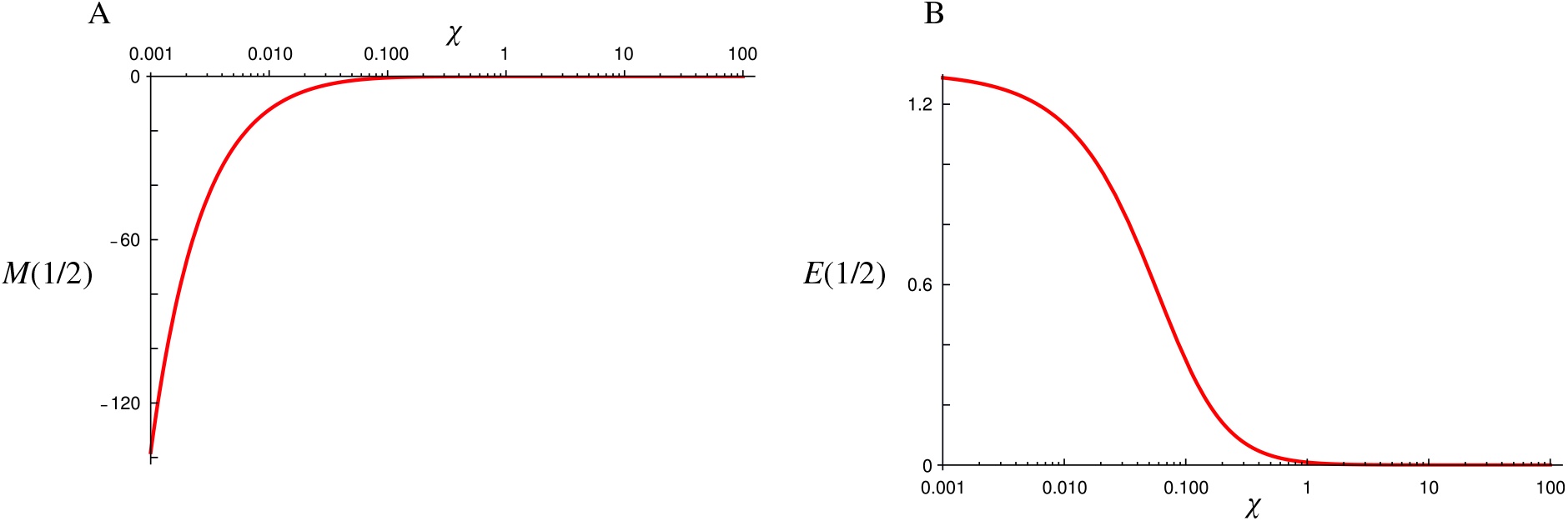
Functions (A) *M*(*x*) and (B) *E*(*x*) evaluated in *x* = 1/2 as functions of the handling time *χ*. The numerical value of both functions significantly changes with *χ*. However, the sign of both *M* and *E* remains constant in the range *χ* ∈ [0.001, 100]. Both functions monotonically approach 0 for *χ* → ∞. Parameter values: *φ* = 10, *σ* = 5, *δ* = 0.8, *γ* = 0.5.

**Figure S2:**
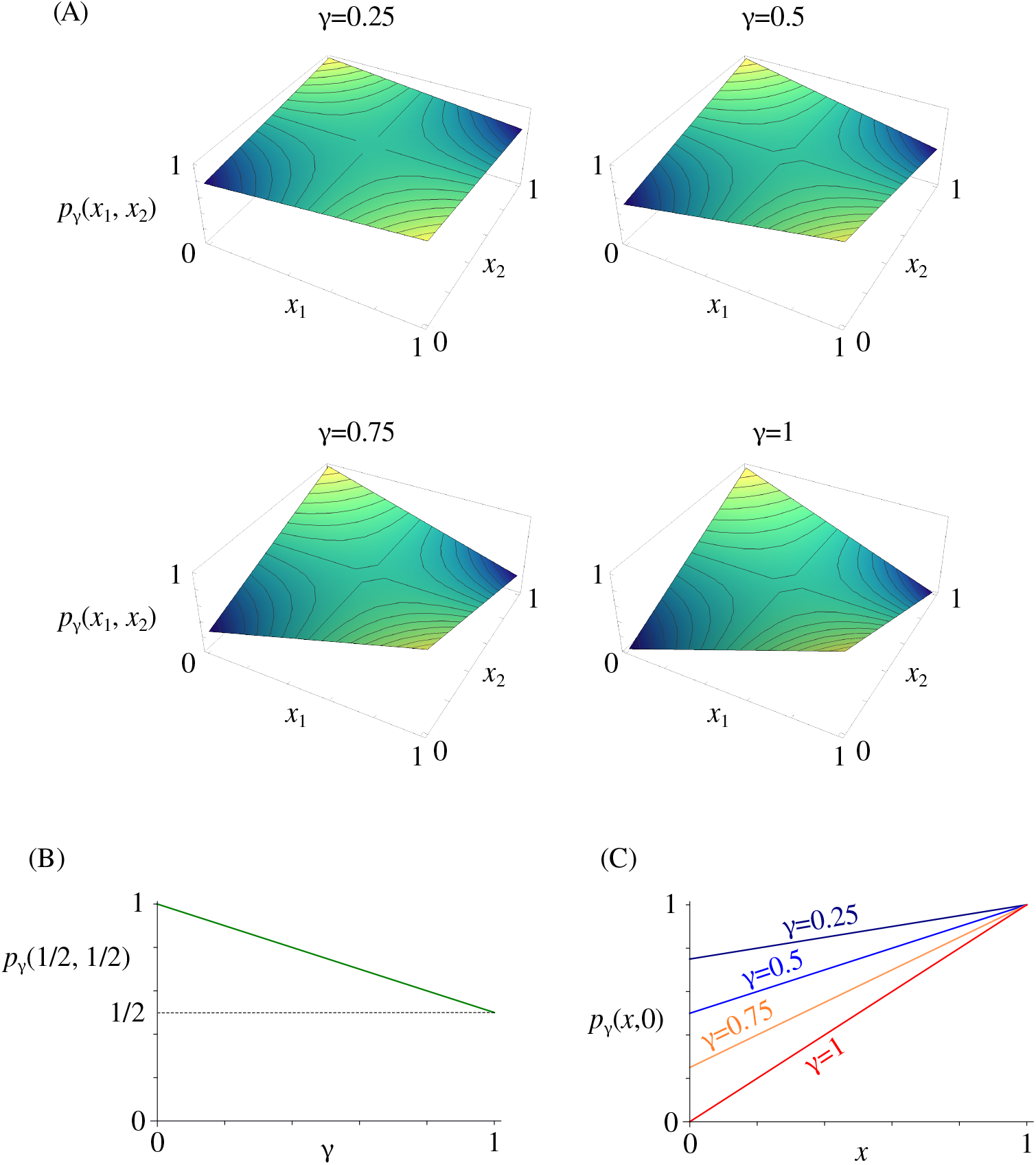
(A) Pollination function *p*_*γ*_(*x*_1_, *x*_2_) for four different numerical values of the efficiency reduction for pollen transfer between flowers of the same type *γ*. (B) Pollination function as a function of *γ* calculated in *x*_1_ = 1/2 = *x*_2_. The value of *p*_*γ*_(*x*_1_, *x*_2_) decreases linearly as *γ* increases. (C) Pollination function as a function of the phenotypic trait *x*_1_ = *x* of a plant when exchanging pollen with a second plant with only L-flowers (*x*_2_ = 0) for four different values of *γ*.

**Figure S3:**
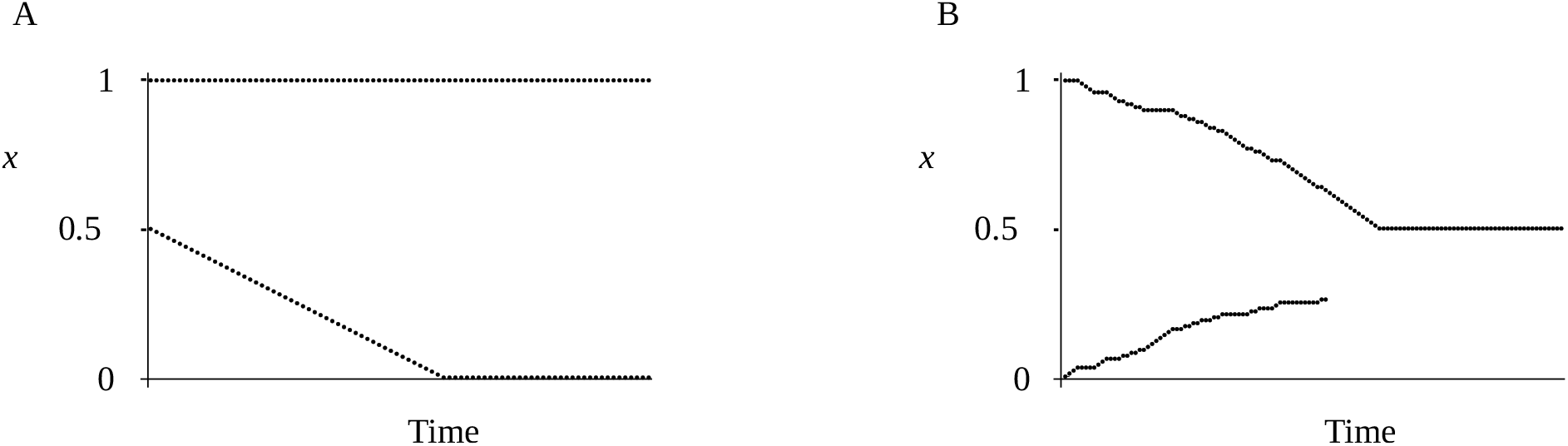
(A) Simulated evolutionary tree for a population initially consisting of trait values *x*_1_ = 1 and *x*_2_ = 1/2. While the trait *x*_1_ is at a fitness peak, the trait value *x*_2_ evolves towards *x* = 0 to reciprocally match the *x*_1_ branch. Parameter values: *φ* = 10, *σ* = 5, *χ* = 1, *δ* = 0.9, *γ* = 0.8. (B) Simulated evolutionary tree of an initially dimorphic population. Due to a decreased reproductive rate caused by, e.g., decline of pollinator service, the phenotypic traits evolve toward *x* = 1/2. Note that, due to stochasticity, the two branches evolve with different speed until one subpopulation outcompetes the other, causing its extinction. Parameter values: *φ* = 5, *σ* = 5, *χ* = 1, *δ* = 0.9, *γ* = 0.8.

**Figure S4:**
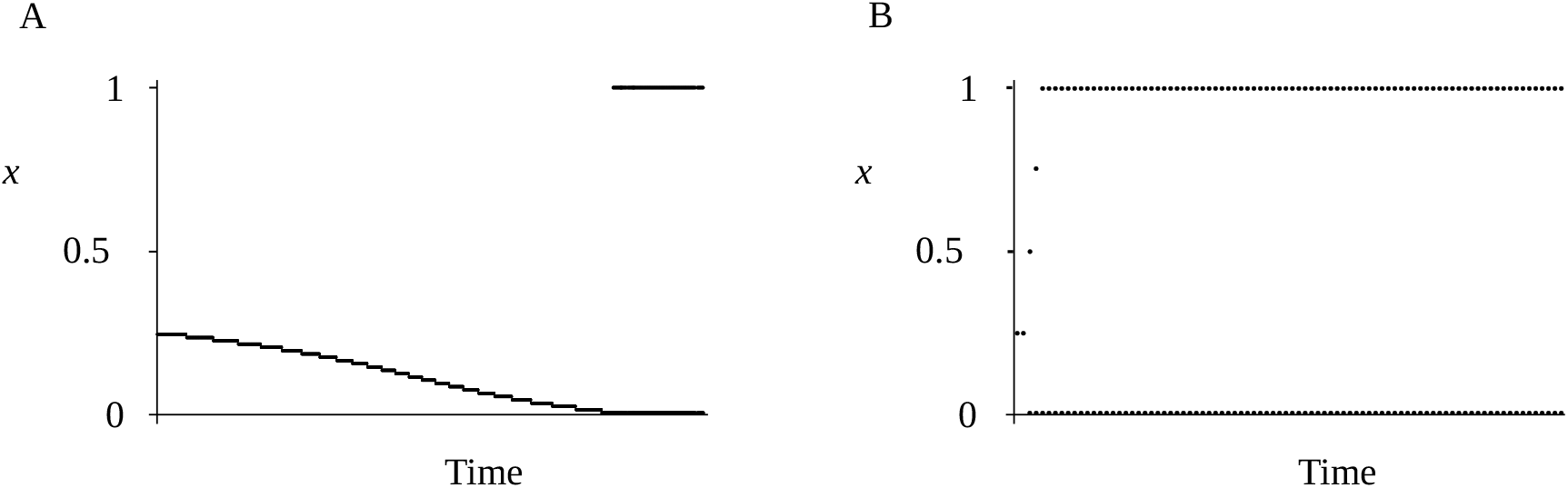
Simulated evolutionary tree in the repeller regime. (A) An initially monomorphic population evolves towards a fixed, uniform direction for stylar deflection *x*_1_ = 0. There, the population is invaded by a second population with *x*_2_ = 1. The now dimorphic population is evolutionary stable. (B) A monomorphic population undergoes evolutionary branching in the repeller regime when the mutation step size is large enough (here, we used mutation step size = 0.25). Parameter values: *φ* = 15, *σ* = 5, *χ* = 1, *δ* = 0.9, *γ* = 0.2.

